# Compositionally Constrained Sites Drive Long Branch Attraction

**DOI:** 10.1101/2022.03.03.482715

**Authors:** Lénárd L. Szánthó, Nicolas Lartillot, Gergely J. Szöllősi, Dominik Schrempf

## Abstract

Accurate phylogenies are fundamental to our understanding of the pattern and process of evolution. Yet, phylogenies at deep evolutionary timescales, with correspondingly long branches, have been fraught with controversy resulting from conflicting estimates from models with varying complexity and goodness of fit. Analyses of historical as well as current empirical datasets, such as alignments including Microsporidia, Nematoda or Platyhelminthes, have demonstrated that inadequate modeling of across-site compositional heterogeneity, which is the result of biochemical constraints that lead to varying patterns of accepted amino acids along sequences, can lead to erroneous topologies that are strongly supported. Unfortunately, models that adequately account for across-site compositional heterogeneity remain computationally challenging or intractable for an increasing fraction of contemporary datasets. Here, we introduce “compositional constraint analysis”, a method to investigate the effect of site-specific constraints on amino acid composition on phylogenetic inference. We show that more constrained sites with lower diversity and less constrained sites with higher diversity exhibit ostensibly conflicting signal under models ignoring across-site compositional heterogeneity that lead to long branch attraction artifacts and demonstrate that more complex models accounting for across-site compositional heterogeneity can ameliorate this bias. We present CAT-PMSF, a pipeline for diagnosing and resolving phylogenetic bias resulting from inadequate modeling of across-site compositional heterogeneity based on the CAT model. CAT-PMSF is robust against long branch attraction in all alignments we have examined. We suggest using CAT-PMSF when convergence of the CAT model cannot be assured. We find evidence that compositionally constrained sites are driving long branch attraction in two metazoan datasets and recover evidence for Porifera as the sister group to all other animals.

---

Understanding the biological foundations of contemporary life on Earth requires detailed knowledge of evolutionary history. The history of speciation events informs us about the appearance of advantageous innovations and the loss of dispensable traits in a continuously changing environment. Consequently, development of phylogenetic models inferring the history of speciation events has continued at an impressive pace during the past decades.

Models of sequence evolution are inevitably simplifications of the complex processes that generate real-life biological sequences. Unfortunately, overly-simplistic models can lead to model misspecification and long branch attraction (LBA; e.g., Felsenstein, 1978; Hendy and Penny, 1989; Zharkikh and Li, 1993; Tateno et al., 1994; Bruno and Halpern, 1999; Ho and Jermiin, 2004; Bergsten, 2005; Brinkmann et al., 2005; Philippe et al., 2011b). LBA is a systematic bias where distantly related lineages are incorrectly inferred to be closely related in reconstructed phylogenies. LBA arises when two lineages appear similar (thus closely related) to one another because they have both undergone a large amount of change, rather than because they are closely related by descent. Probabilistic substitution models (e.g., Jukes and Cantor, 1969) can account for multiple substitutions per site and as a result reduce LBA (Felsenstein, 1973; but see Farris, 1999) compared to methods that do not correct for multiple substitutions.

However, probabilistic substitution models may still yield biased estimates if they do not adequately describe the evolutionary processes. For example, model violation may occur due to improper description of the heterogeneity of the substitution process across sites (Yang, 1993), across branches (Tuffley and Steel, 1998), or more fine-grained modulations through time (heterotachy and heteropecilly: Philippe and Lopez, 2001; Roure and Philippe, 2011), all of which have motivated a lot of work (e.g., Galtier, 2001; Huelsenbeck, 2002; Lopez et al., 2002; Kolaczkowski and Thornton, 2004; Philippe et al., 2005b; Lockhart et al., 2006; Zhou et al., 2007; Lartillot et al., 2009; Jayaswal et al., 2011; Crotty et al., 2020). In the present work, the focus is on heterogeneity across sites.

Historically, the dichotomy between invariable and variable sites was the first to be considered: inference with models ignoring invariant sites can be severely biased, and conversely, just accounting for a proportion of invariable sites leads to substantial improvement (Shoemaker and Fitch, 1989; Adachi and Hasegawa, 1995a; Lockhart et al., 1996). In a more quantitative spirit, models accounting for rate heterogeneity across sites (i.e., allowing for slower or faster evolution at different sites) ameliorate LBA in some cases (Kuhner and Felsenstein, 1994; Philippe et al., 2011b). Heterogeneity of the process is not restricted to rates, however. Empirically (e.g. Jayaswal et al., 2014), amino acid and nucleotide composition can vary both across sites (i.e., columns of an alignment) and across lineages or taxa (i.e., rows of an alignment).

The interaction between sites in a protein sequence is sometimes referred to as “intramolecular epistasis” (Noor et al., 2012). In particular, we will consider preferences for specific types of nucleotides or amino acids at homologous positions in alignments of such data, reflected in compositional variation between columns of an alignment. Such “compositional heterogeneity across sites” is a result of site specific selection constraints, for example, deeply buried positions in folded proteins tend to have more interactions and are correspondingly more constrained compared to positions on the surface, which have fewer interactions and are less constrained (Koshi and Goldstein, 1995; Yeh et al., 2014; Jimenez et al., 2018).

Compositional constraints at a site reflect selection on multiple timescales. Interactions between sites induced by structural and functional constraints maintained by natural selection constrain the site-specific amino acid composition over long evolutionary timescales. Compositional constraints may also result from short term fluctuations in site-specific amino acid preferences resulting from changes at integrating sites, which are relaxed as compensatory substitutions occur (Pollock et al., 2012).

Amino acid and nucleotide composition, however, varies not only across sites, but also across branches and time, reflected in compositional variation between rows of an alignment. Such shifts in compositional preference are often driven by environmental changes and life history traits, for example by differences in temperature driving proteome-wide amino acid composition across prokaryotes (Boussau et al., 2008). Accounting for across branch and across time changes in composition requires using nonstationary substitution models (Foster et al., 1997; Jermiin et al., 2004). In this paper, we assume stationarity of the evolutionary process across the branches of the tree and focus on modeling compositional heterogeneity across sites in the alignment.

Our primary motivation is that models ignoring across-site compositional heterogeneity are prone to LBA because they underestimate the probability of convergent substitutions at compositionally constrained sites (Lartillot and Philippe, 2004). The probability of independent substitutions to the same state depends on the number of acceptable amino acids, which differs across sites. Models ignoring across-site compositional heterogeneity pool all sites, and ignore the variation of the evolutionary process across sites. Indeed, analyses of a series of datasets exhibiting previously contentious evolutionary relationships provide evidence that ignoring across-site compositional heterogeneity can lead to LBA (Phillips et al., 2004; Brinkmann et al., 2005; Philippe et al., 2005b,a; Delsuc et al., 2006; Lartillot et al., 2007; Philippe et al., 2009, 2011a; Brown et al., 2013; Ryan et al., 2013; Cannon et al., 2016; Simion et al., 2017).

There is accumulating evidence that accounting for across-site heterogeneities is key to an accurate reconstruction of deep evolutionary relationships. The classic approach to modeling such heterogeneities in the phylogenetic inference process are mixture models that combine substitution models specifically tailored to the evolutionary processes observed in the data. In order to model across-site rate heterogeneity, we use a mixture of substitution models with the same relative but different absolute substitution rates (e.g., Yang, 1993; Kalyaanamoorthy et al., 2017).

Modeling across-site compositional heterogeneity, however, requires constructing a process describing the evolution of sites subject to different compositional constraints. To do so for time-reversible substitution models, we can leverage the separation of substitution rates into the product of (1) symmetric exchangeabilities, describing differences in rates of exchange between pairs of states, and (2) stationary frequencies of the target states (e.g., Whelan and Goldman, 2001). Figuratively, the stationary frequencies of the target states correspond to the probabilities of sampling the target states after waiting for an infinitely long time. Any time-reversible substitution model is fully specified by a set of symmetric exchangeabilities and the set of stationary frequencies (sometimes also referred to as a profile or a stationary distribution, e.g., Lartillot and Philippe (2004)). Assuming biochemical constraints primarily affect site-specific amino acid preferences in the long term, across-site compositional heterogeneity can be accounted for by composing a number of substitution models sharing a single set of exchangeabilities but differing in their stationary distributions (distribution mixture models, sometimes also referred to as profile mixture models; Quang et al., 2008; Schrempf et al., 2020). We note that distribution mixture models are usually augmented with a model accounting for across-site rate heterogeneity (e.g., Yang, 1993). Thus, distribution mixture models should be an adequate choice even when the evolutionary rate correlates with amino acid composition (Gowri-Shankar and Rattray, 2005).

We can distinguish between general distribution mixture models estimated from curated training databases, and distribution mixture models directly estimated from the datasets at hand. For example, Wang et al. (2008) directly estimate mixture model components using principal component analysis. Susko et al. (2018) use a composite likelihood approach and additional methods such as taxon weighing. In contrast, the rationale behind providing and using general mixture models is the assumption that the underlying evolutionary processes share universal features. Quang et al. (2008) use the expectation maximization algorithm to infer general mixture models consisting of 10, 20,…, 60 components (C10, C20,…, C60, collectively CXX models). Schrempf et al. (2020) used a clustering approach together with different compositional transformations to provide a set of general mixture models, termed universal distribution mixtures (UDM), with the number of components ranging from four up to several thousand. They also provide the clustering method EDCluster to infer dataset-specific distribution mixture models.

Finite mixtures can be used in the context of maximum likelihood (ML) phylogenetic inference. However, statistical analyses of model fit and investigation of known cases of LBA indicate that a large number of components are necessary for robustness against LBA (Schrempf et al., 2020) which is computationally expensive especially in terms of random-access memory.

Bayesian approaches can more easily accommodate richer mixtures. Nonparametric Bayesian methods do not require explicit specification of the number of mixture components nor their stationary distributions. In particular, the CAT model (Lartillot and Philippe, 2004) uses a Dirichlet process prior to approximate an arbitrary mixture of stationary distributions across sites. The CAT model was shown to be better fitting and less prone to LBA than site-homogeneous models for a series of classic datasets including Nematoda and Platy-helminthes (Lartillot et al., 2007) as well as in phylogenomic analyses of the tree of life (Williams et al., 2020). The impediment of nonparametric Bayesian methods, and specifically the CAT model, is that it compounds two computationally challenging, but separately tractable problems, the nonparametric inference of the underlying distribution across sites and the exploration of tree space. The combination of these two problems is challenging and can lead to convergence problems.

In all cases, mixture modeling approaches accounting for across-site compositional heterogeneity are complex and require considerable computational resources (e.g., Whelan and Halanych, 2016). In order to reduce computational cost, Wang et al. (2018) proposed a two-step approximation. First, site-specific stationary distributions are estimated using a reference mixture model and a fixed guide topology. We note that the choice of the guide topology affects the estimation of the stationary distributions, and that a suboptimal guide topology may bias the results. Second, a tree is inferred using the fixed stationary distributions obtained in the first step. Thereby, runtime is reduced while robustness against LBA is improved compared to using the reference mixture model alone. In particular, the site-specific stationary distributions are set to the posterior mean site frequencies (PMSF) of the reference mixture model. As a result, the phylogenetic accuracy of the PMSF approach is inherently limited by how well the reference mixture model captures across-site compositional heterogeneity. There is no reason to restrict the use of the PMSF approach to empirical mixture models: any random-effect model meant to account for pattern heterogeneity could in principle be used here as a reference mixture model for computing the posterior means of the site-specific stationary distributions.

In this work we follow a multistep procedure similar to the PMSF model and address two points: First, on the computational side, we extend the PMSF approach by using the CAT model instead of an empirical profile mixture model as the reference model for computing the profiles. Importantly, the CAT model is used under a fixed guide topology. Thus, the problem of simultaneous inference of both the site-specific stationary distributions and the tree is reduced to a search of site-specific stationary distributions, and tree branch lengths only. We termed our approach CAT-PMSF.

Second, we investigate the contribution of sites with different degrees of compositional constraints to LBA. In particular, we test to what extent more severely constrained sites exhibit bias towards LBA trees under models that do not adequately account for across-site compositional heterogeneity. We employ “compositional constraint analysis”, which examines phylogenetic signal for alternative topologies as a function of per site compositional diversity measured by the effective number of amino acids (see Materials and Methods).

Examining simulated alignments as well as classic empirical datasets including Microsporidia (Brinkmann et al., 2005), Nematoda, and Platyhelminthes (Philippe et al., 2005a), we find conflicting phylogenetic signal across sites with different degrees of compositional constraints. Based on these results we apply compositional constraint analysis to datasets analyzed previously by Ryan et al. (2013) and Simion et al. (2017), aiming to resolve the early diversification of animal lineages.

## Materials and Methods

In the following, we use the terms *site-homogeneous* and *site-heterogeneous* when referring to models ignoring and accounting for across-site compositional heterogeneity, respectively. Further, we use the term *tree* to denote a directed acyclic graph with node labels and branch lengths, in which exactly one branch connects any two nodes. We use the term *topology* to denote a tree without information about branch lengths but with node labels. We specify an evolutionary model with exchangeabilities, EX, and across-site compositional heterogeneity model, ASCH as EX+ASCH. All discussed evolutionary models used for simulations as well as inferences implicitly use discrete gamma rate heterogeneity with four components. We add a flag +PMSF to denote usage of the posterior mean site frequency model (Wang et al., 2018).

### Effective Number of Amino Acids

Given a distribution *π* = (*π_A_*, *π_R_*,…, *π_V_*) of amino acid frequencies, we seek a simpler measure *K*_eff_(*π*) in the closed interval [1, 20] that indicates the effective number of different amino acids used. We refer to this number as the “effective number of amino acids”, and denote it as *K*_eff_ (Schrempf et al., 2020). Lartillot et al. (2007) and Pollock et al. (2012) refer to the same concept, calling it the “effective size of the amino acid alphabet” or the “biochemical diversity”.

There are at least two conceptual platforms to compute *K*_eff_: (1) Homoplasy, that is, the probability of sampling the same amino acid twice, and (2) the Shannon entropy. Lartillot et al. (2007) and Pollock et al. (2012) use the Shannon entropy. Schrempf et al. (2020) examine both concepts and did not observe significant differences between the two definitions of *K*_eff_. Here, we prefer to compute *K*_eff_ using homoplasy because the probability of LBA directly depends on the probability of homoplasy. Briefly, if the phylogenetic model underestimates the probability of homoplasy in the alignment, sequence similarities may be wrongly attributed to a potential “close evolutionary distance”, and not to potential “random similarity because of homoplasy”.

In particular, the probability of homoplasy is

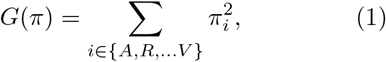

and the effective number of amino acids is the inverse

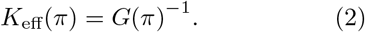

*K*_eff_ is a convenient measure because it ranges from 1.0, when one amino is used exclusively, to 20.0 for a uniform distribution. Further, we can apply *K*_eff_ to the distribution of amino acid frequencies at a given site. In this case, *K*_eff_ denotes the “number of amino acids used at a given site”. Finally, we note that Wright (1990) also uses the concept of homoplasy (which population geneticists call “homozygosity”) to define a more elaborate measure of the “effective number of codons used in a gene” (see also Fuglsang, 2006).

### CAT-PMSF

Figure 1 shows an overview of the CAT-PMSF pipeline. The input to CAT-PMSF is an alignment. The output of the CAT-PMSF pipeline is a tree robust to LBA.

**Figure 1:**
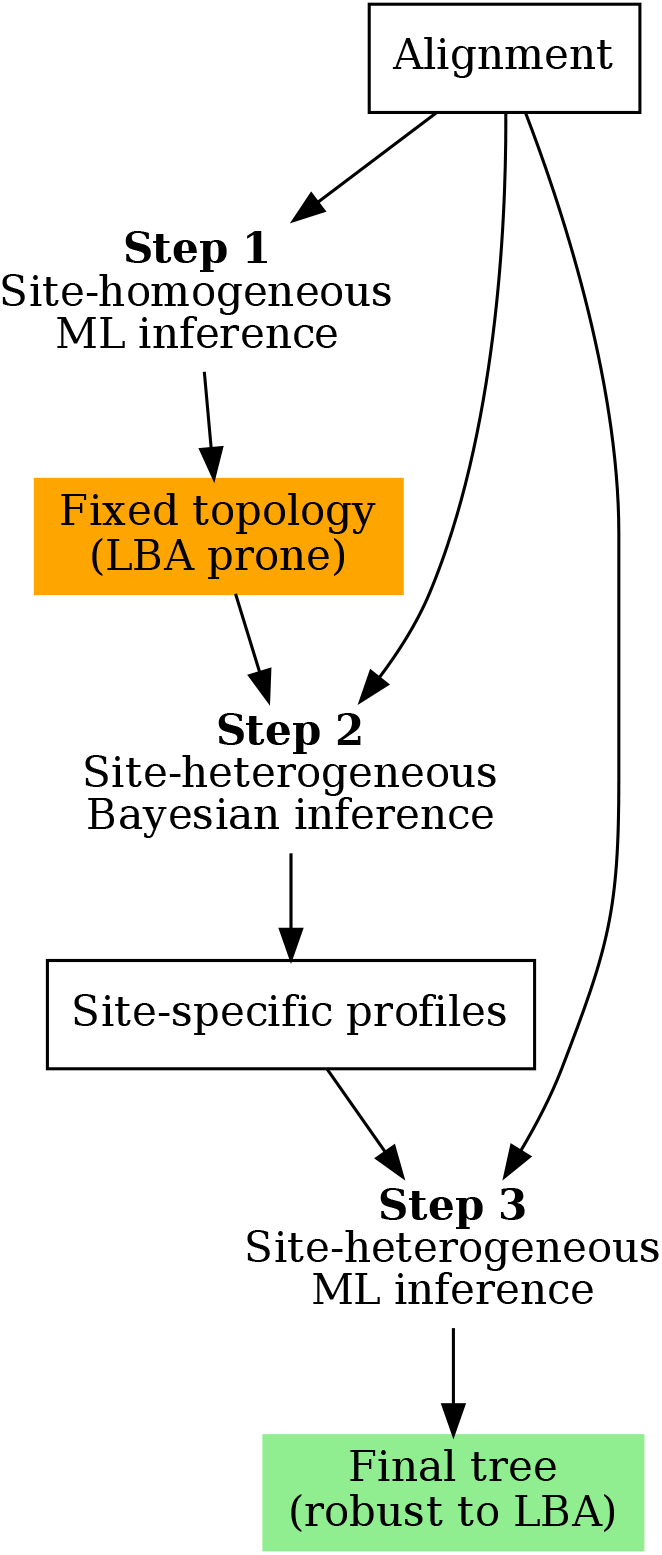
The CAT-PMSF pipeline. (1) Apply a site-homogeneous maximum likelihood (ML) model (LG+G4; Yang, 1993; Le and Gascuel, 2008) with IQ-TREE 2 (Minh et al., 2020). The obtained tree may still suffer from long branch attraction. (2) Fix the topology of this tree in a site-heterogeneous inference with the Bayesian CAT model (Lartillot and Philippe, 2004) and extract the posterior mean site-specific stationary distributions of amino acids. (3) Estimate a tree robust to long branch attraction with the obtained site-specific stationary distributions in IQ-TREE 2.

#### Step 1

Use a site-homogeneous model to infer a ML tree with IQ-TREE 2 (Minh et al., 2020). Specifically, we used LG exchangeabilities (Le and Gascuel, 2008), the empirical stationary distribution of amino acids, and a discrete gamma rate model (Yang, 1993) with 4 categories (LG+F+G4 in IQ-TREE 2 terminology).

#### Step 2

Use the topology of the obtained tree, which may be biased by LBA, in a subsequent Bayesian analysis with the CAT model (Lartillot and Philippe, 2004) in PhyloBayes (Lartillot et al., 2013). Analogous to the PMSF approach, call this the “guide topology”. Fix the guide topology during this step of the CAT-PMSF pipeline to reduce the computational requirements of the CAT model. Then, extract the posterior mean site-specific stationary distributions of amino acids. In our analyses, we either used Poisson (Felsenstein, 1973; Nei, 1987), LG (Le and Gascuel, 2008) or GTR (Tavaré, 1986) exchangeabilities, and a discrete gamma rate model with 4 categories. We ran two Markov chains until either the effective sample size of all parameters was above 100, or after visual inspection with Tracer (Rambaut et al., 2018) indicated convergence. Due to computational constraints the Markov chains involving the Simion et al. (2017) and Ryan et al. (2013) alignments do not reach an estimated sample size of 100 for each parameters, for detail see Tables from S16 to S28. For the GTR model, we also extracted the posterior mean exchangeabilities from the results of PhyloBayes. All scripts are available in the Dryad Digital Repository: https://doi.org/10.5061/dryad.g79cnp5rh and at https://github.com/drenal/cat-pmsf-paper.

#### Step 3

Use the custom site-specific stationary distributions in IQ-TREE 2. To this end, use capabilities of IQ-TREE 2 implemented as part of the PMSF method (Wang et al., 2018). The PMSF method has two steps. First, infer the site-specific stationary distributions. Second, use the inferred site-specific stationary distributions for phylogenetic inference. Here, we use the second step of the PMSF method together with the custom site-specific stationary distributions obtained in Step 2 of the CAT-PMSF pipeline.

### Simulations

In order to assess the accuracy of CAT-PMSF, we simulated alignments of 10000 amino acids under a distribution mixture model (Schrempf et al., 2020). We used Poisson exchangeabilities (Felsenstein, 1973; Nei, 1987) and a discrete gamma rate model (Yang, 1993) with 4 categories with shape parameter *α* = 0.78. The distribution mixture model has site-specific stationary distributions. For each site, we sampled a random distribution from a universal set of distributions (Schrempf et al., 2020) obtained from the HOGENOM (Dufayard et al., 2005) and HSSP (Schneider et al., 1997) databases.

We used Felsenstein-type trees with four leaves (insets in top row of Figure 2; Felsenstein, 1978). Felsenstein-type trees exhibit two long branches separated by a short internal branch. The quartet trees had different branch length proportions between short (*q*) and long (*p*) branches. We fixed *q* to 0.1, and changed *p* between 0.1 and 2.0. We stored the randomly sampled site-specific stationary distributions used for the simulation. For the simulations we used the ELynx suite (https://github.com/dschrempf/elynx). The scripts and the simulated data are available in the Dryad Digital Repository: https://doi.org/10.5061/dryad.g79cnp5rh and at https://github.com/drenal/cat-pmsf-paper.

**Figure 2:**
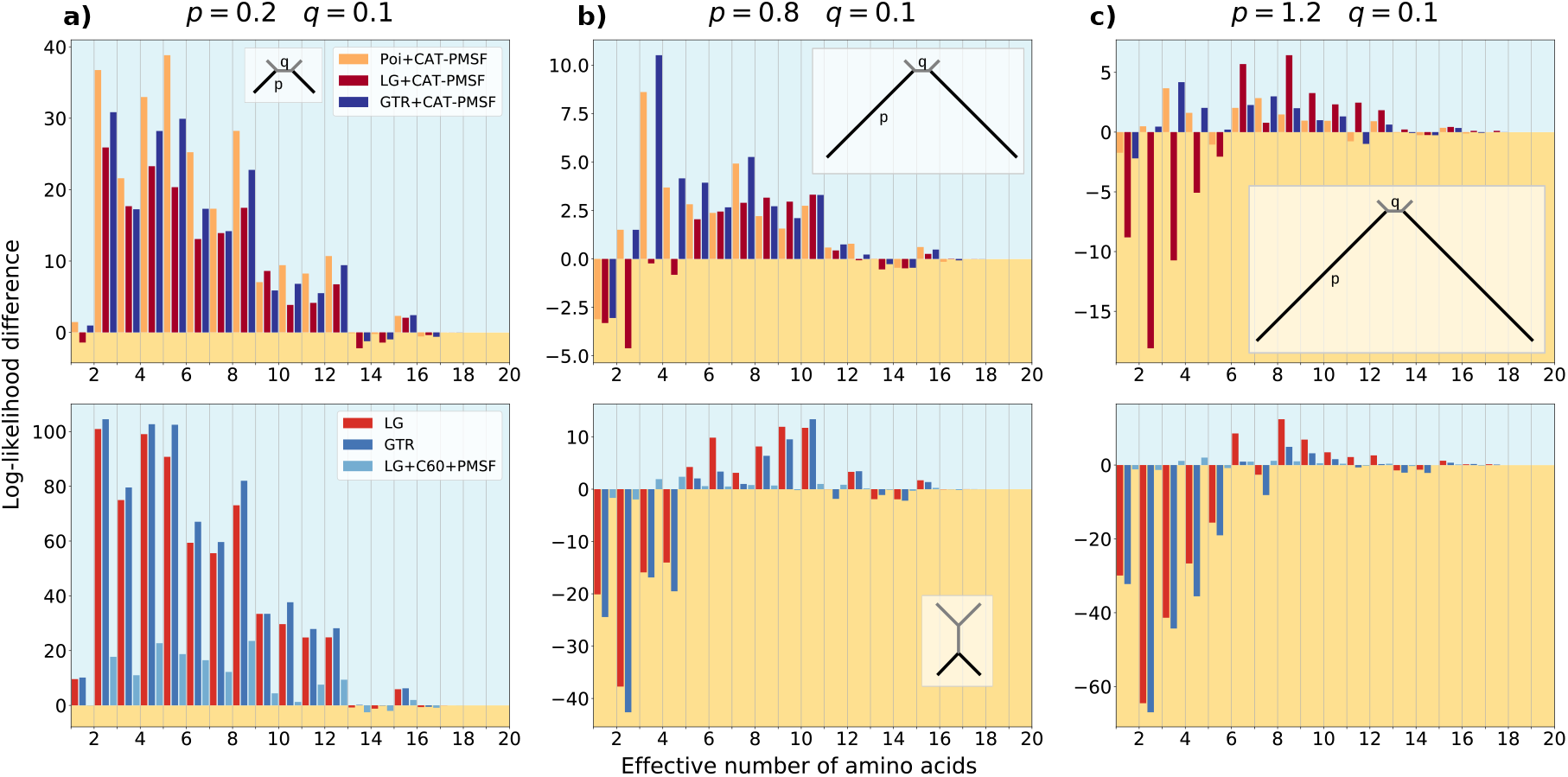
Highly constrained sites drive long branch attraction artifacts in the Felsenstein zone. We simulated amino acid alignments with 10 000 sites exhibiting across-site compositional heterogeneity (Schrempf et al., 2020) along Felsenstein-type trees (insets in top row; Felsenstein, 1978) with different branch lengths *q* = 0.1, and *p* = 0.3, 0.8, and 1.2 from (a) to (c). We performed analyses with CAT-PMSF, the Poisson (Felsenstein, 1973; Nei, 1987), the LG (Le and Gascuel, 2008) and the GTR (Tavaré, 1986) models constrained to the correct topology as well as to an incorrect topology (inset in bottom row; Farris, 1999) with IQ-TREE 2 (Minh et al., 2020). The site-specific loglikelihood differences ΔlogL between the maximum likelihood trees of the two competing topologies binned according to the site-specific effective number of amino acids *K_eff_* are shown. A positive value (blue background) indicates support for the true topology, a negative value (yellow background) indicates support for the incorrect topology exhibiting long branch attraction. The LG, and GTR models incorrectly infer Farris-type trees if *p* ≥ 0.8.

### Compositional Constraint Analysis

We calculated the site-specific likelihood differences between two analyses constrained to two different topologies (-g flag in IQ-TREE 2), let us denote them topology *A*, and *B*, respectively. For example, in the simulation study, we chose topology B such that the inferred tree was of the Felsenstein-type (insets in top row of Figure 2). Felsenstein-type trees exhibit a short internal branch separating two long extant branches. We chose topology A such that the inferred tree was of the Farris-type (inset in bottom row of Figure 2(b)). Farris-type trees have two long extant branches that merge before joining a short internal branch. For a comparison of Felsenstein-type and Farris-type trees, see for example Leuchtenberger et al. (2020). For each site *i*, we calculated the loglikelihood difference as

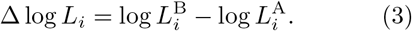

A positive value of Δlog *L_i_* indicates that site i supports topology B. A negative value indicates support for topology A.

We ordered and binned the sites according to their *K*_eff_ values and summed the site-specific loglikelihood differences within each bin. We chose 20 bins because there are 20 amino acids, but different bin sizes may be used (Fig. S13, S14, and S15 available in the Supplementary Material which can be found in the Dryad Digital Repository: https://doi.org/10.5061/dryad.g79cnp5rh). For the simulation study, we used the actual *K*_eff_ values of the sampled amino acid profiles during the simulation. For the analyses of empirical datasets, we used the *K*_eff_ values calculated from the site-specific stationary distribution obtained in Step 2 of the CAT-PMSF pipeline. Taxa in the insets of Figure 3 and Figure 4 are represented using Phylopic (http://phylopic.org/), the silhouette for Microsporidia is based on Tosoni et al. (2002, Figure 7).

**Figure 3:**
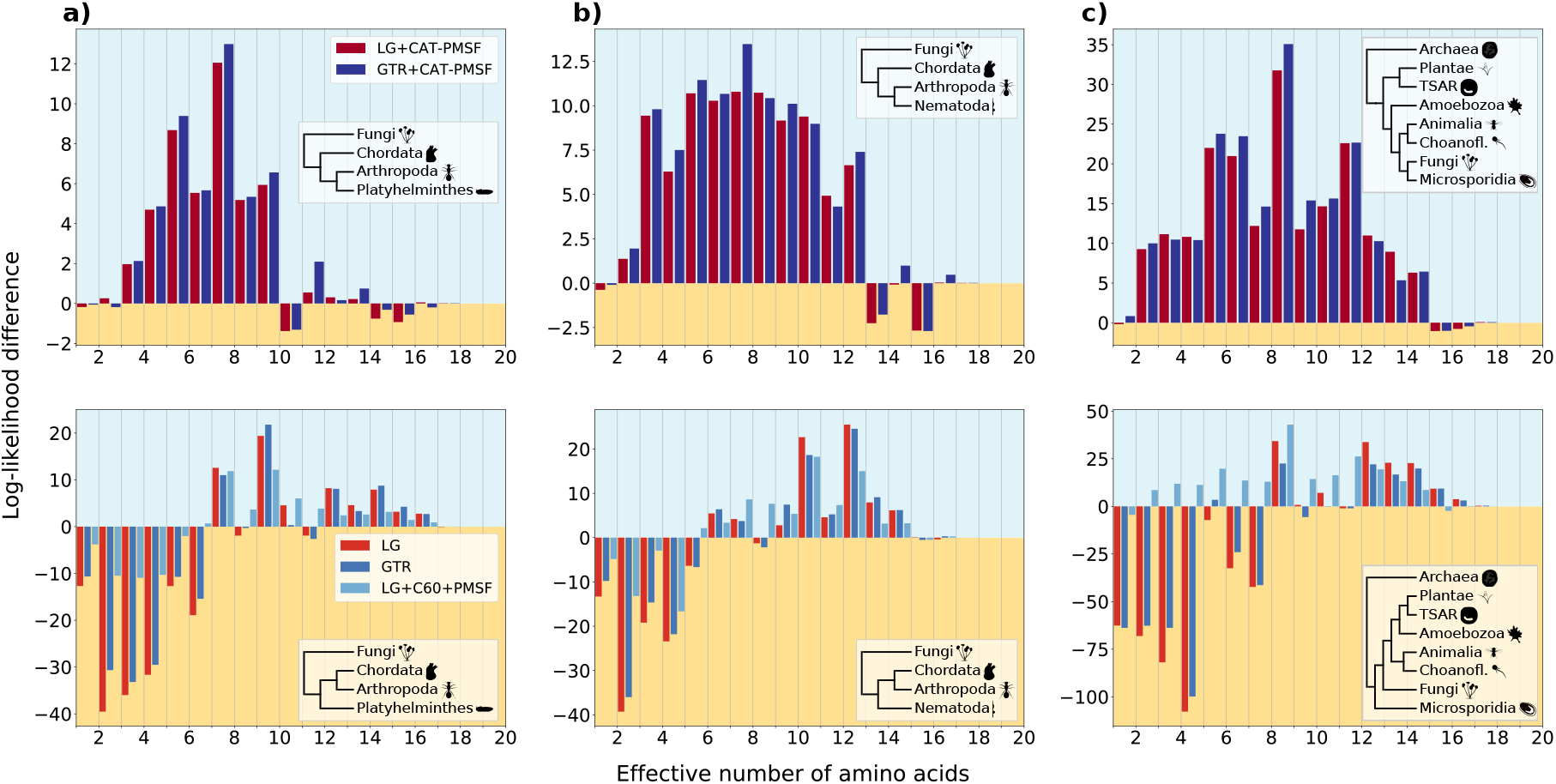
Highly constrained sites explain classic examples of long branch attraction. We analyzed three empirical datasets including (a) Platyhelminthes and (b) Nematoda (Philippe et al., 2005a), and (c) Microsporidia (Brinkmann et al., 2005). We performed analyses with CAT-PMSF, the LG (Le and Gascuel, 2008), the GTR (Tavaré, 1986), and the LG+C60+PMSF (Quang et al., 2008; Wang et al., 2018) models constrained to either one of two competing topologies (insets in top versus bottom rows) with IQ-TREE 2 (Minh et al., 2020). The site-specific log-likelihood differences ΔlogL between the LBA-prone and non-LBA-prone topologies binned according to the site-specific effective number of amino acids *K*_eff_ estimated by PhyloBayes (Lartillot and Philippe, 2004) are shown. A positive value (blue background) indicates support for the now accepted topology, a negative value (yellow background) indicates support for the topology prone to long branch attraction. Site-homogeneous models infer the wrong topology for all three datasets.

**Figure 4:**
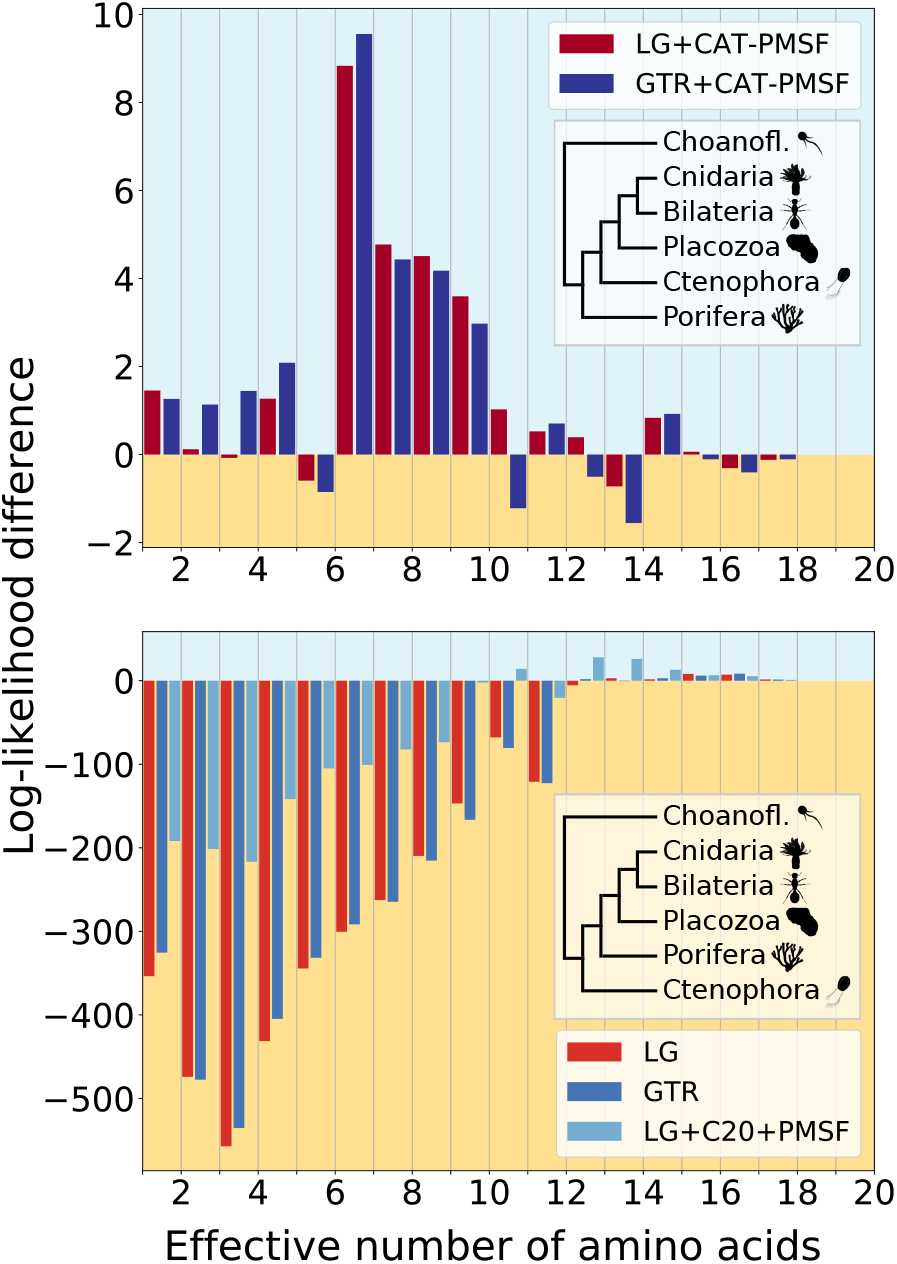
CAT-PMSF shows consistent sig-nal for Porifera as the sister group to all other animals on the alignment from Simion et al. (2017). We performed analyses with CAT-PMSF, the LG (Le and Gascuel, 2008), the GTR (Tavaré, 1986), and the LG+C20+PMSF (Quang et al., 2008; Wang et al., 2018) models constrained to either one of two competing topologies (insets in top versus bottom rows) with IQ-TREE 2 (Minh et al., 2020) on the alignment from Simion et al. (2017). The site-specific log-likelihood differences ΔlogL between the maximum likelihood trees of the two competing topologies binned according to the site-specific effective number of amino acids *K*_eff_ estimated by PhyloBayes (Lartillot and Philippe, 2004) are shown. Site-homogeneous models and the site-heterogeneous LG+C20+PMSF model show inconsistent signal between more versus less constrained sites and favor Ctenophora at the animal root. CAT-PMSF favors Porifera at the animal root, although this result is only significant when using the closest outgroup exclusively.

### Dataset involving Platyhelminthes and Nema-toda

Philippe et al. (2005a) address a well-known LBA artifact concerning the placement of Platyhelminthes and Nematoda on the tree of Bilateria. Lartillot et al. (2007) revisit the same dataset and provide two reduced, and overlapping alignments which contain 37 species for Nematoda and 32 species for Platyhelminthes, respectively. Both alignments have a length of 35371 amino acids. Figure 3 (a) and (b), S1 and S2 show simplified and complete species trees, respectively.

### Dataset involving Microsporidia

The dataset provided by Brinkmann et al. (2005) comprises 40 species with 24294 amino acids. It contains an archaean outgroup and eukaryotic taxa. Of particular interest are the Microsporidia, a group of unicellular parasites which lack mitochondria and instead possess mitosomes. Microsporidia evolve fast, and site-homogeneous methods fail to correctly classify them. Application of site-heterogeneous methods confirms that Microsporidia are the closest sister species of Fungi (Brinkmann et al., 2005). For these reasons, the dataset containing Microsporidia is ideal as a proof of concept for CAT-PMSF. Figure 3 (c) and S3 show simplified and complete species trees, respectively.

### Metazoan Datasets

The placement of Ctenophora on the tree of Metazoa is still a matter of debate. We apply CAT-PMSF to two datasets. First, the alignment provided by Ryan et al. (2013) contains 61 species with 88384 amino acids. Second, the alignment provided by Simion et al. (2017) contains 97 species with 401632 amino acids. The complete set of outgroups comprises 2 Filasterea, 5 Ichthyosporea, and 18 Choanoflagellatea. The Choanoflagellatea are the closest outgroup. Figure 4 shows results obtained from a reduced alignment in which we retained only the Choanoflagellatea. The reduced alignment yields 90 species. Figure 4, and Figure S4 show simplified and complete species trees, respectively.

### Compositional Heterogeneity across Branches

CAT-PMSF assumes homogeneity of the evolutionary process across the branches of the tree. The matched-pairs test of symmetry (Ababneh et al., 2006) tests for compositional heterogeneity between two sequences. Homo v2.1 (https://github.com/lsjermiin/Homo.v2.1) performs this test for all pairs of sequences in an alignment. We applied Homo v2.1 to the simulated as well as empirical alignments.

## Results

In brief, CAT-PMSF comprises three steps: (1) estimate a guide topology using a site-homogeneous model, (2) estimate site-specific stationary distributions with the CAT model in PhyloBayes (Lartillot and Philippe, 2004) using the guide topology, and (3) phylogenetic inference in a ML framework with a distribution mixture model sharing one set of exchangeabilities, and using the obtained site-specific stationary distributions (see Materials and Methods).

### Simulation Study

We assessed and compared the accuracy of CAT-PMSF with other site-homogeneous and site-heterogeneous models. To this end, we simulated amino acid sequence alignments with a length of 10 000 sites along Felsenstein-type quartet trees (insets in top row of Figure 2; Felsenstein, 1978). We used uniform exchangeabilities (Poisson; Felsenstein, 1973) and an across-site compositional heterogeneity model with site-specific stationary distributions based on a UDM model (see Material and Methods; Schrempf et al., 2020). We set the branch length of the short branch *q* to 0.1, and varied the length of the long branch *p* between 0.1 and 2.0. As expected, using Homo v2.1, we did not detect evidence for compositional heterogeneity across sequences in the simulated alignments (Supplementary Section Compositional heterogeneity across sequences).

The true topology was not recovered with site-homogeneous models when *p* ≥ 0.8. Instead, a Farris-type (or LBA) tree is recovered (Fig. S7, S8, and Tables S1, S2). Figure 2 shows the results of the compositional constraint analysis for different values of *p* = 0.2, 0.8 and 1.2, contrasting the site-wise log-likelihood differences between the ML trees constrained to the true (Felsenstein-type) and incorrect (Farris-type) topologies exhibiting LBA (see Materials and Methods). Figures S7, and S8 show results for other values of p. We binned sites according to the effective number of amino acids (*K*_eff_, see Materials and Methods) used by the respective site profiles. Lower values of *K*_eff_ correspond to sites under stronger compositional constraints. Compositional constraint analysis compares per site phylogenetic signal for two alternative topologies as a function of *K*_eff_. Here, positive log-likelihood differences indicate support for the true topology. Conversely, negative values indicate support for the alternative topology exhibiting LBA. In the absence of model misspecification, we expect consistent phylogenetic signal (i.e., support for either one or the other topology) across sites, and independent of the true value of *K*_eff_.

At odds with this expectation, site-homogeneous evolutionary models exhibit conflicting phylogenetic signal between sites with lower and higher *K*_eff_ values (Fig. 2, S7 and S8). For site-homogeneous evolutionary models, more constrained sites with a lower value of *K*_eff_ exhibit bias towards the incorrect topology exhibiting LBA. For *p* ≥ 0.8, the bias outweighs the correct signal of less constrained sites with high values of *K*_eff_, and the incorrect topology has higher support than the true topology. In contrast, the site-heterogeneous LG+C60+PMSF and Poisson+CAT-PMSF models show consistent support for the true topology.

To ascertain the statistical significance of the compositional constraint analysis we calculated Pearson’s correlation coefficients, as well as Spearman’s and Kendall’s rank correlation coefficients and associated p-values between the loglikelihood differences and the site-specific *K*_eff_ values (Supplementary Section Measuring correlation between *K*_eff_ and site-specific log-likelihood difference; Tables S3, S5 and S7). For the Pearson’s correlation coefficient, site-homogeneous models exhibit large and significant correlation for *p* ≥ 0.8, whereas the log-likelihood differences and *K*_eff_ values of site-heterogeneous models are not correlated.

Approximately unbiased (AU) tests (Shimodaira, 2002) of ML trees inferred by the GTR+CAT-PMSF model constrained to the two alternative topologies reject the incorrect topology exhibiting LBA in favor of the true topology for *p* < 1.5 (Table S9). AU tests of the Poisson+CAT-PMSF model show similar results in that the topology exhibiting LBA is rejected for *p* < 1.6 (Table S10). The LG+CAT-PMSF model only rejects the topology exhibiting LBA for *p* < 0.8, and favors the incorrect topology for *p* > 1.0 (Table S10).

Finally, we note that site-heterogeneous models with GTR exchangeabilities perform well if *p* ≤ 1.4. This is an interesting observation, because the simulations used the Poisson model with uniform exchangeabilities, and both Phylobayes and IQ-TREE 2 use LG exchangeabilities as starting values when inferring GTR exchangeabilities (e.g., see the results of the GTR+CAT-PMSF model in Figure 2c). This indicates, that the inference of exchangeabilities has converged well if *p* ≤ 1. 4. In contrast, model misspecification by fixing the exchangeabilities to the ones of the LG model indeed leads to inconsistent signal similar to the one obtained with site-homogeneous models when *p* ≥ 0.8 (e.g., see the results of the LG+CAT-PMSF model in Figure 2c). The Poisson+CAT-PMSF model is even more accurate when using the true topology or the true site-specific stationary distributions (Fig. S7 and S8). In all cases excluding the LG+CAT-PMSF and GTR+CAT-PMSF models for large p, the site-heterogeneous models infer the true topology (Fig. S7 and S8).

### Applications to empirical Data

Similar to the simulation study above, for empirical alignments the site-specific stationary distributions obtained in Step 2 of the CAT-PMSF pipeline can be used to quantify the strength of compositional constraint measured by *K*_eff_ and perform compositional constraint analysis. Figure 3 shows results for three datasets exhibiting classic LBA artifacts when we use site-homogeneous models for inference: The placement of Platyhelminthes and Nematoda (Philippe et al., 2005a), as well as the placement of Microsporidia (Brinkmann et al., 2005; Lartillot et al., 2007). For all three datasets, Homo v2.1 reported some level of compositional heterogeneity across sequences (see Materials and Methods and Supplementary Section Compositional heterogeneity across sequences). Figures S9 and S11 show the cumulative site-specific log-likelihood differences and Figures S13, S14 and S15 provide results for alternative bin sizes.

For site-homogeneous models, the site-specific log-likelihood differences between the ML trees constrained to the two competing topologies (insets of Figure 3; top vs. bottom) show conflicting phylogenetic signal between more or less constrained sites. The bias towards the topologies exhibiting LBA artifacts of more constrained sites outweighs the signal of less constrained sites in all three datasets.

The site-heterogeneous LG+C60+PMSF model shows reduced, but still apparent conflict compared to site-homogeneous models and the LG+C10+PMSF model (Fig. S9). For Platyhelminthes, the bias is strong enough that the total likelihood across all sites is higher for the LBA topology, while for the datasets involving Nematoda and Microsporidia, the LG+C10+PMSF model recover the correct topology, albeit with reduced support. In general, the results of the LG+C10+PMSF and LG+C60+PMSF models are consistent with the observation (Schrempf et al., 2020) that increasing the number of mixture model components of the CXX models decreases the bias introduced by more constrained sites. Pearson correlation coefficients are greater for site-homogeneous models than for models LG+C10+PMSF and LG+C60+PMSF, but significant for each of these (Tables S4, S6 and S8).

In contrast, CAT-PMSF exhibits consistent signal towards the assumed-to-be-correct topologies across all sites and datasets with no significant correlation between log-likelihood difference and site-specific *K*_eff_ value (Table S4). The ML trees inferred by CAT-PMSF are consistent with the accepted phylogenetic relationships and AU tests confirm the rejection of trees with LBA topologies (Tables S12-S14).

### The phylogenetic Position of Ctenophora

Finally, we used CAT-PMSF on two metazoan datasets (Ryan et al., 2013; Simion et al., 2017) to investigate early evolutionary relationships on the animal tree of life. It is currently a matter of debate whether sponges (Porifera) or comb jellies (Ctenophora) are the sister group to all other animals (e.g., Kapli and Telford, 2020; Li et al., 2021). We refer to the competing hypotheses as Porifera-sister and Ctenophora-sister, respectively.

Compositional constraint analysis under site-homogeneous models, as well as combinations of PMSF and site-heterogeneous mixture models with 20 and 60 components exhibit patterns of conflicting phylogenetic signal for sites with different degrees of compositional constraints (Fig. 4, S10 and S12) for both the alignments from Simion et al. (2017) and Ryan et al. (2013). The conflicting signal is consistent with LBA driving the placement of Ctenophora as the first animal group to emerge (Table S4).

Under site-homogeneous models, sites with *K*_eff_ values up to approximately 10 – 12 exhibit strong preference for Ctenophora-sister (Fig. 4 and S17). Sites with higher *K*_eff_ values, however, switch their preference toward Porifera-sister. In contrast, under the CAT-PMSF models the Simion et al. (2017) dataset exhibits consistent phylogenetic signal (Table S3) favoring a Porifera-sister topology and rejecting the Ctenophora-sister topology (AU test p-values between 3.1 × 10^-4^ and 7.7 × 10^-4^; Table S15) with the closest outgroup, Choanoflagellatea (Fig. S4a, Figure S10). For the alignment published by Ryan et al. (2013), the total log-likelihood difference of CAT-PMSF between the two hypotheses is marginal at only 0.8, suggesting a lack of resolution in this dataset. None of the models we investigated exhibit consistent phylogenetic signal across sites with different degrees of compositional constraints.

## Discussion

We introduce CAT-PMSF, a method for phylogenetic inference from alignments exhibiting across-site compositional heterogeneity. The CAT-PMSF pipeline uses the site-specific amino acid preferences estimated by a nonparametric Bayesian approach in the context of a downstream ML analysis. Doing so combines the benefits of both approaches: a more accurate inference of the patterns across sites with a computationally more efficient and more reproducible inference of the tree topology. In addition to phylogenetic inference, the CAT-PMSF pipeline can also be used to investigate the consistency of phylogenetic signal for sites under different degrees of compositional constraints. Compositional constraint analyses on both simulated and empirical datasets exhibiting across-site compositional heterogeneity show that site-homogeneous and some site-heterogeneous mixture models indeed have inconsistent signal which contributes to topological bias and LBA artifacts.

In the simulation study (Fig. 2), site-homogeneous models favored the incorrect topology when the length *p* of the terminal branches was long enough. By separating the contribution of sites as a function of compositional constraint, we demonstrated that more constrained sites drive the bias leading to LBA.

The threshold *K*_eff_ values separating sites supporting the correct topology and sites supporting the LBA topology depended on the length *p* of the terminal branches: The longer the terminal branches, the higher the threshold *K*_eff_ value. We expect this observation holds more generally.

In our simulations, support of site-homogeneous models shifted from the true topology towards the incorrect topology when increasing *p* to and above 0.8. In this case, sites with *K*_eff_ values above that threshold failed to compensate for the bias introduced by sites with *K*_eff_ values below the threshold. We observed no bias when using site-heterogeneous models such as Poisson+CAT-PMSF (Fig. 2). Although we expect such a result, it is satisfying that inferences of Poisson+CAT-PMSF lack bias even for large values of *p* ≥ 1.2 (Fig. S7 and S8).

We discovered bias towards one of the topologies in simulation studies because we know the true parameters and trees. Bias is harder to detect in analyses of empirical data. Compositional constraint analysis detects conflicting signal between more and less constrained sites. Detection of such inconsistencies is a strong indicator for bias: Knowing the stationary distribution of a site alone should not provide us with information about the favored evolutionary history. In mathematical terms, the log-likelihood difference of a site between two hypotheses should be conditionally independent given the stationary distribution of that site. In contrast, we expect the signal obtained from more and less constrained sites be consistent up to random statistical error.

In our analyses of empirical data we observed strong inconsistencies between more and less constrained sites for site-homogeneous models and hardly any inconsistencies when using CAT-PMSF (Fig. 3). Pearson correlation coefficients and p-values confirm this observation across a wide range of simulated and empirical datasets (Tables S3, and S4).

The results are more nuanced for the alignments involving Ctenophora. In the case of site-homogeneous models, we observe the value of *K*_eff_ correlates strongly with the log-likelihood difference between the two competing topologies (Fig. 4). Moreover, for the dataset provided by Simion et al. (2017), CAT-PMSF supports Porifera-sister — similar to the results reported by the original authors, who applied the CAT model to sub-sampled alignments comprising 100 000 sites. The support of CAT-PMSF for Porifera-sister is significant when we use the closest outgroups exclusively (Table S15). If we add more distant outgroups, the results are less conclusive (Table S15). LBA provides an explanation for this observation: outgroups more distant to the ingroup (i.e., the species of interest) attract outgroups closer to the ingroup (Hendy and Penny, 1989; Adachi and Hasegawa, 1995b) thereby increasing the distance between the ingroup and the whole outgroup. Consequently, the longer basal branch increases bias due to LBA for branches leading to the most recent common ancestor of the ingroup. In particular, the elongated basal branch of animals increases bias due to LBA for branches leading to the metazoan root.

We interpret these findings as a confirmation for sponges being the sister group to all other animals (dataset of Simion et al., 2017), and believe that the inconclusive results obtained from the dataset of Ryan et al. (2013) reflect a lack of phylogenetic resolution. Irrespective of the final evolutionary history of Metazoa, our results add important evidence that ignoring across-site compositional heterogeneity leads to LBA (Phillips et al., 2004).

Compositional constraint analysis seeks to decompose the log likelihood contributions of more and less constrained sites. We decided to measure how constrained sites are by computing the effective number of amino acids per site based on the concept of homoplasy, because it is inherently related to LBA: A model that under-estimates the probability of homoplasy will incorrectly attribute sequence similarities due to homoplasy to a putative close evolutionary distance. Other measures of how constrained sites are or any monotonic transformation of the function computing *K*_eff_ could be used. For example, the Shannon entropy can be used to calculate a slightly different measure of the effective number of amino acids, albeit with similar characteristics (Schrempf et al., 2020). To reiterate, compositional constraint analysis is useful, because we do not expect that the topological preference depends on how constrained sites are. Indeed, if all sites have been produced under the same evolutionary history, then they should agree on the preferred topology (or, possibly, abstain, e.g., if they are constant), and this, even if they otherwise differ in other aspects of the evolutionary process (such as the biochemical constraints). In particular, we expect that the sign of the site-specific log-likelihood difference does not change between more or less constrained sites. However, this is exactly what we observe for almost all inferences when using site-homogeneous models and even for some inferences with site-heterogeneous models (e.g., Figures 2–4). More quantitatively, the correlation coefficients between *K*_eff_ and site-specific log-likelihood difference tend to be stronger for site-homogeneous models than for site-heterogeneous models (Tables S3, S5 and S7).

We also note that CAT-PMSF assumes homogeneity of the evolutionary process across branches of the tree. The simulation study conforms to this assumption. Tests for across-branch homogeneity were less conclusive for the empirical datasets (Supplementary Section Compositional heterogeneity across sequences) than for the simulation study. If the evolutionary process is stationary and homogeneous, the CAT model should perform well in estimating site-specific amino acid compositions. Even if the evolutionary process is non-stationary or heterogeneous, the site-specific amino acid compositions inferred by the CAT model will capture the spectrum of compositions attained at least somewhere on the tree. In this case, the inferred compositions will not be stationary in the mathematical sense, but still should have a positive impact on the detection of genuine convergent evolution. Ideally, we should model across-site and across-branch compositional heterogeneity for amino acid sequences in a combined way. For example, there has been work in this direction on nucleotide sequences (e.g. Jayaswal et al., 2014). In the case of amino acids, one could begin with a simulation study testing if (dis)-similarity in amino acid composition influences evolutionary distances or even the topology estimated by CAT-PMSF or other methods accounting for across-site compositional heterogeneity.

The results of CAT-PMSF are conservative because the CAT model estimates the site-specific stationary distributions using guide topologies prone to LBA artifacts. That is, the guide topologies are obtained with site-homogeneous models. Even so, CAT-PMSF correctly infers the true trees in the simulation study (Table S1), and trees that we are convinced to be free from LBA artifacts in the analyses comprising empirical datasets (Fig. S3, S2, S1, S6, S5 and S4). This observation justifies the usage of site-homogeneous models in Step 1 of the CAT-PMSF pipeline.

In the simulation study, we observe bias towards Farris-type trees when using site-homogeneous models, and no bias or reduced bias when using Poisson+CAT-PMSF or GTR+CAT-PMSF. However, the reduced bias comes at a cost: the absolute values of the log-likelihood differences are greater for site-homogeneous models than for site-heterogeneous models, even though site-heterogeneous models are more parameter rich.

In general, site-homogeneous models show conflicting signal between more and less constrained sites, but we observe hardly any such inconsistencies when using CAT-PMSF. In any case, even when the signal across sites is consistent, evidence obtained from highly constrained sites should be examined carefully, especially when highly constrained sites weigh more heavily than less constrained sites. We are convinced that inconsistencies between more and less constrained sites are a strong indicator for the presence of LBA.

Li et al. (2021) argue that only the most parameter rich models favor Porifera-sister, and so Porifera-sister is not a likely scenario. In contrast, Schrempf et al. (2020) report that statistical tests favor models using more stationary distributions. This point is confirmed here, where we see that CXX models, in spite of being generally more robust against LBA than site-homogeneous models, may still be insufficient and result in conflicting signal (Fig. S8, S9 and S10). In practice, many sites may have evolved under different conditions, so we can not expect all sites to share a universal stationary distribution. In fact, we do not even expect stationarity. In our opinion, we should analyze data using complex models and decide about which parameters are necessary to grasp the complexities of evolution. With CAT-PMSF we further explored this path. The CAT-PMSF method uses site-specific stationary distributions and therefore is a parameter-rich model.

In comparison, the site-specific posterior mean stationary distributions of the classical PMSF approach are a superposition of a finite set of stationary distributions of the underlying mixture model. Consequently, the *K*_eff_ values of the site-specific stationary distributions of the classical PMSF approach must be equal to or larger than the lowest *K*_eff_ value of the stationary distributions of the underlying mixture model. Further, we expect even the richest distribution mixture models do not offer adequate variability of components with stationary distributions exhibiting low *K*_eff_ values. For example, there are twenty different stationary distributions with *K*_eff_ values close to 1.0, 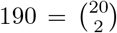 stationary distributions with *K*_eff_ values close to 2.0, and so on.

Finally, the speed benefit of CAT-PMSF originates from fixing the topology during the Bayesian analysis with the CAT model. Of course, estimating the site-specific stationary distributions is still by far the most time-consuming step. In the future, we aim to design improved methods estimating site-specific stationary distributions. Specifically, we are thinking about methods based on machine learning such as AlphaFold (Jumper et al., 2021).

In conclusion, compositional constraint analyses show evidence for a potential LBA caused by model misspecification, and an argument that careful model choice as well as validation is important in phylogenetic inference. We also propose a method, CAT-PMSF, with the potential to produce more accurate phylogenetic estimates using a site heterogeneous, but branch homogeneous, substitution process.

## Acknowledgments

The authors thank Tom A. Williams, Edu Ocaña-Pallarès and László G. Nagy for constructive criticism of the manuscript.

## Supplementary Material

Supplementary material, including data files and/or online-only appendices, can be found in the Dryad Digital Repository: https://doi.org/10.5061/dryad.g79cnp5rh.

