## Supplemental Material for "Compositionally Constrained Sites Drive Long Branch Attraction"

### Compositionally Constrained Sites Drive Long Branch Attraction. Supplementary Material.

March 1, 2023

S5 AU tests 31

S6 Filtering out distant outgroups helps reduce long branch attraction 37

S7 Results for another Metazoa dataset 39

S8 Compositional heterogeneity across sequences 40

S9 Convergence measures of Phylobayes analyses 43

#### S1 Commands to run the CAT-PMSF pipeline

Figure 1 shows the required steps to apply the CAT-PMSF pipeline on a phylogenetic dataset. Although other implementations exist and may be used for the various steps, our analyses were conducted solely using IQ-TREE 2 (Minh et al., 2020) and PhyloBayes (Lartillot and Philippe, 2004). Step 1 can be done via IQ-TREE 2 (Minh et al., 2020) on an alignment preferably in PHYLIP (Felsenstein, 1989) format, because Phylobayes (Lartillot and Philippe, 2004) expects that as input type in Step 2.

```
iqtree2 -s alignment.phylip -m LG+F+G4
```

This will output a maximum likelihood (ML) tree file called `alignment.phylip.treefile`. Step 2 uses PhyloBayes MPI (Lartillot and Philippe, 2004). Before running Step 2, benchmarks can be performed to choose the best number of parallel tasks the MPI program should use.

```
mpirun -np 48 pb_mpi -cat -gtr -d alignment.phylip -T alignment.phylip.treefile cat_gtr_alignment_chain1
mpirun -np 48 pb_mpi -cat -gtr -d alignment.phylip -T alignment.phylip.treefile cat_gtr_alignment_chain2
```

To assess convergence of the two chains, the guidelines for *acceptable* runs as stated in the PhyloBayes MPI documentation (https://github.com/bayesiancook/pbmpi/blob/master/pb_mpiManual1.8.pdf) are used. That is, the entries of the `tracecomp` command should have effective sample sizes above 50 and relative differences below 0.3.

```
# Phylobayes MPI 1.8 or Phylobayes 4.1c
tracecomp -x 1000 1 cat_gtr_alignment_chain1 cat_gtr_alignment_chain2
# prior to Phylobayes MPI 1.8 or prior Phylobayes 4.1c the burnin argument may only support one value
tracecomp -x 1000 cat_gtr_alignment_chain1 cat_gtr_alignment_chain2
```

The `-x` switch defines the number of burn-in iterations (`1000`) and the sampling frequency (1).

An example output for a successful run for the Nematoda dataset with model CAT+GTR+G4 is shown below.

```
# tracecomp -x 1000 gtr_nematode_icc_chain1 gtr_nematode_icc_chain2
    name                effsize         rel_diff
    loglik              540             0.15339
    length              5339            0.250924
    alpha               4724            0.182099
    Nmode               1292            0.0321017
    statent             301             0.00237055
    statalpha           419             0.180082
    rrent               342             0.130712
    rrmean              6179            0.00141907
```

To export the site-specific stationary distributions from a converged PhyloBayes MPI run, one can use the `readpb_mpi` utility.

```
readpb_mpi -ss -x 1000 1 cat_gtr_alignment_chain1
```

This command will produce the file `cat_gtr_alignment_chain1.siteprofiles`.

In case PhyloBayes was run with GTR exchangeabilities, the inferred exchangeability matrix also has to be exported and supplied in Step 3 of the CAT-PMSF pipeline:

```
readpb_mpi -rr -x 1000 1 cat_gtr_alignment_chain1
```

The output file is `cat_gtr_alignment_chain1.meanrr`.

The custom script converting outputs of PhyloBayes MPI to IQ-TREE 2 compatible formats can be found in the GitHub repository in the Dryad Digital Repository: https://doi.org/10.5061/dryad.g79cnp5rh and at https://github.com/drenal/cat-pmsf-paper. The usage is as follows:

```
python3 convert_site-dists_v2.py cat_gtr_alignment_chain1.siteprofiles
```

This will produce `cat_gtr_alignment_chain1.sitefreq`.

Similarly a script is provided in the GitHub repository in the Dryad Digital Repository: https://doi.org/10.5061/dryad.g79cnp5rh and at https://github.com/drenal/cat-pmsf-paper which converts the exchangeabilities:

```
python3 convert-exchangeabilities.py cat_gtr_alignment_chain1.meanrr
```

This will create `cat_gtr_alignment_chain1.exchangeabilities`

Step 3 of the CAT-PMSF pipeline consists of supplying IQ-TREE 2 (Minh et al., 2020) the posterior mean site-specific stationary distributions of amino acids (and the posterior mean exchangeabilities in case of the GTR model), and running IQ-TREE 2 on the alignment:

```
# for custom exchangeabilities:
iqtree2 -s alignment.phylip -m cat_gtr_alignment_chain1.exchangeabilities+G4 -fs
↪  cat_gtr_alignment_chain1.sitefreq --prefix alignment.phylip.gtrcatpmsf
# for predefined exchangeabilities, e.g. LG:
iqtree2 -s alignment.phylip -m LG+G4 -fs cat_lg_alignment_chain1.sitefreq --prefix
↪  alignment.phylip.lgcatpmsf
```

This will produce the `alignment.phylip.gtrcatpmsf.treefile` or `alignment.phylip.lgcatpmsf.treefile` ML tree inferred using the specified custom model.

#### S2 Trees for empirical datasets

The insets of Figures 3 and 4 show reduced topologies. In this section, we provide the complete ML trees of the datasets inferred by the CAT-PMSF method at Step 3 and the site-homogeneous method at Step 1. Due to the high variance of the branch lengths, we provide the topology with the real branch length denoted in labels above the branches. Red dots at the nodes of the trees signal differences between the topology inferred in Step 1 and 3 of the CAT-PMSF pipeline and correspond to the Robinson-Foulds metric (Robinson and Foulds, 1981). The full trees for Philippe et al.'s Nematoda and Platyhelminthes and Nematoda datasets in Figures S1 and S2 respectively, the full trees for Brinkmann et al.'s Microsporidia dataset are in Figure S3. The full trees for Simion et al.'s Metazoa dataset are in Figures S5 and S4, and the full trees of the other Metazoa dataset from Ryan et al. are in Figure S6.

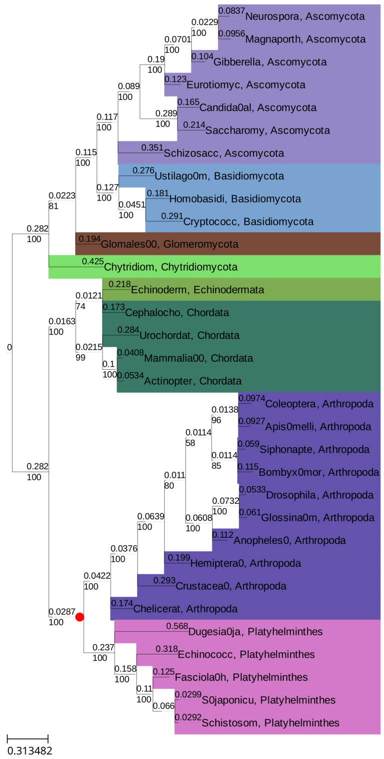

(a) ML tree of Step 3

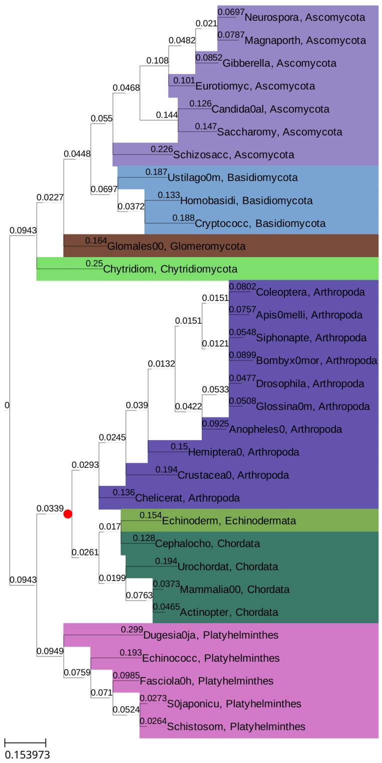

(b) ML tree of Step 1

Figure S1: **Detailed trees for Philippe et al.'s Platyhelminthes dataset.** Maximum likelihood topologies inferred by the site-heterogeneous GTR+CAT-PMSF model (a), and the site-homogeneous LG model (b) of the Philippe et al. (2005) Platyhelminthes dataset representing 32 species. Color codes are the same for species of the same taxa. Red dots mark differences between topologies (a) and (b) and correspond to the Robinson-Foulds metric (Robinson and Foulds, 1981). Values above branches represent inferred branch lengths, values below the branches represent bootstrap support. On the inset topology in Figure 3a, Chordata include Echinodermata; and Chytridiomycota, Glomeromycota, Basidiomycota and Ascomycota form Fungi.

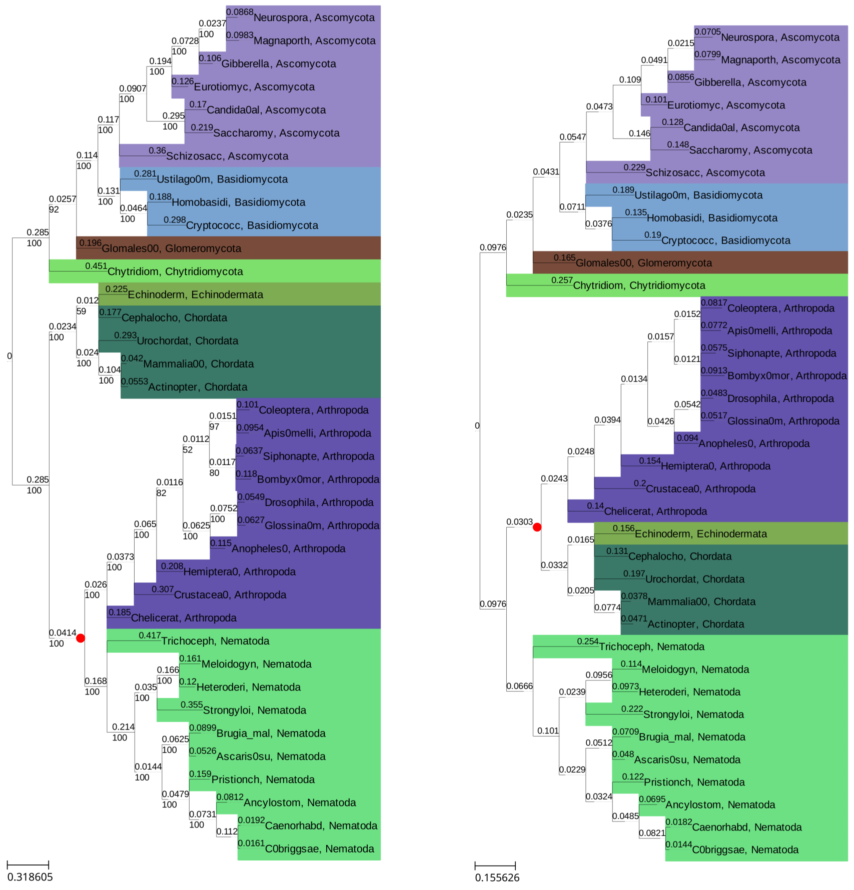

(a) ML tree of Step 3 (b) ML tree of Step 1

Figure S2: **Detailed trees for Philippe et al.'s Nematoda dataset.** Maximum likelihood topologies inferred by the site-heterogeneous GTR+CAT-PMSF model (a), and the site-homogeneous LG model (b) of the Philippe et al. (2005) Nematoda dataset representing 37 species. Color codes are the same for species of the same taxa. Red dots mark the differences between topologies (a) and (b) and correspond to the Robinson-Foulds metric (Robinson and Foulds, 1981). Values above branches represent inferred branch lengths, values below branches represent bootstrap support. On the inset topology in Figure 3b, Chordata include Echinodermata; and Chytridiomycota, Glomeromycota, Basidiomycota and Ascomycota form Fungi.

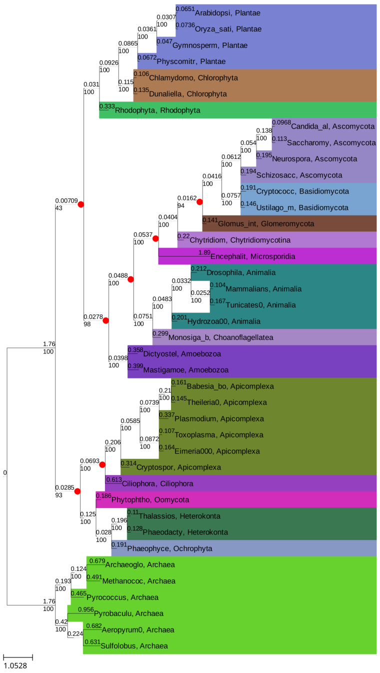

(a) ML tree of Step 3

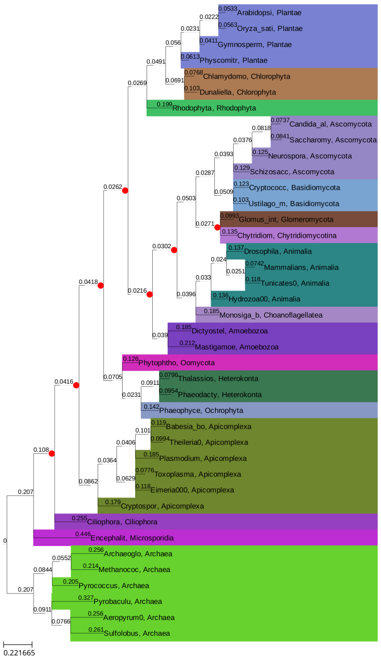

(b) ML tree of Step 1

Figure S3: **Detailed trees for the Brinkmann et al. dataset.** Maximum likelihood topologies inferred by the site-heterogeneous GTR+CAT-PMSF model (a), and the site-homogeneous LG model (b) of the Brinkmann et al. (2005) dataset representing 40 species. Color codes are the same for species of the same taxa. Red dots mark differences between topologies (a) and (b) and correspond to the Robinson-Foulds metric (Robinson and Foulds, 1981). Values above branches represent inferred branch lengths, values below branches represent bootstrap support. The inset topology in Figure 3c collects the phyla Apicomplexa, Ciliphora, Oomycota, Heterokonta and Ochrophyla under the supergroup TSAR; whereas there is no distinction between Plantea, Chlorophyta and Rhodophyta. Ascomycota, Basidiomycota, Glomeromycota and Chytridiomycotina form the Fungi group.

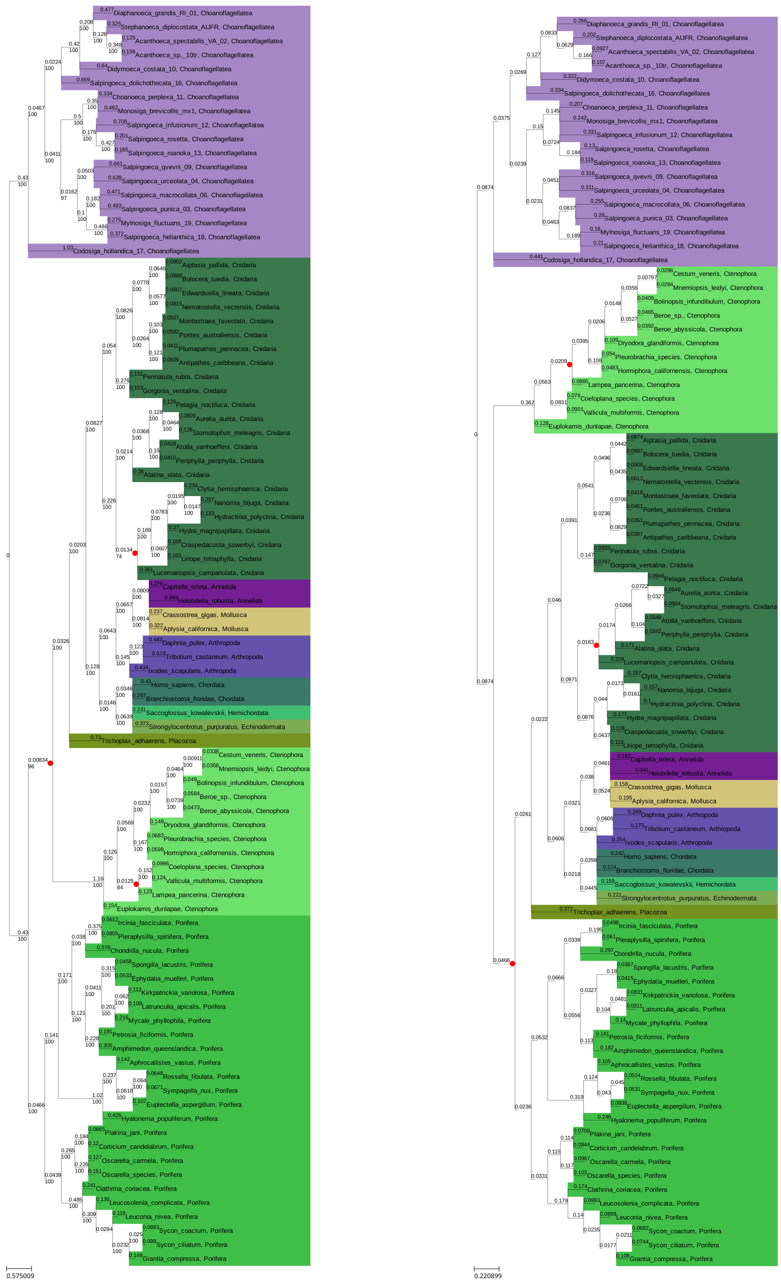

(a) ML tree of Step 3 (b) ML tree of Step 1

Figure S4: **Detailed trees for Simion et al.'s reduced dataset.** Maximum likelihood topologies inferred by the site-heterogeneous GTR+CAT-PMSF model (a), and the site-homogeneous LG model (b) of the reduced Simion et al. (2017) dataset representing 90 species. Color codes are the same for species of the same taxa. Red dots mark the differences between trees (a) and (b) and correspond to the Robinson-Foulds metric (Robinson and Foulds, 1981). Values above branches represent inferred branch lengths, values below branches represent bootstrap support.

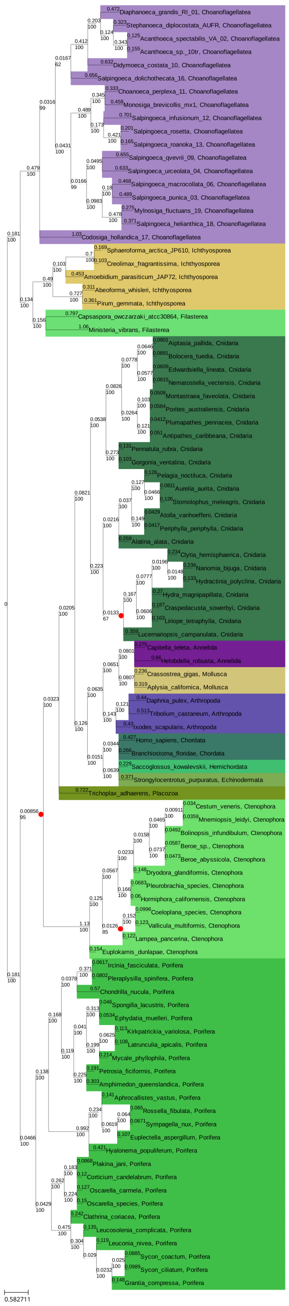

(a) ML tree of Step 3

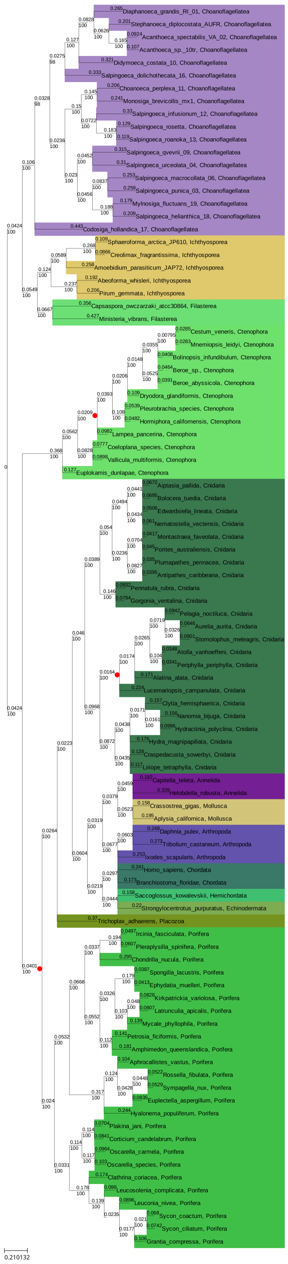

(b) ML tree of Step 1

Figure S5: **Detailed trees for Simion et al.'s full dataset.** Maximum likelihood topologies inferred by the site-heterogeneous GTR+CAT-PMSF model (a), and the site-homogeneous LG model (b) of the Simion et al. (2017) dataset representing 97 species. Color codes are the same for species of the same taxa. Red dots mark differences between topologies (a) and (b) and correspond to the Robinson-Foulds metric (Robinson and Foulds, 1981). Values above branches represent inferred branch lengths, values below branches represent bootstrap support.

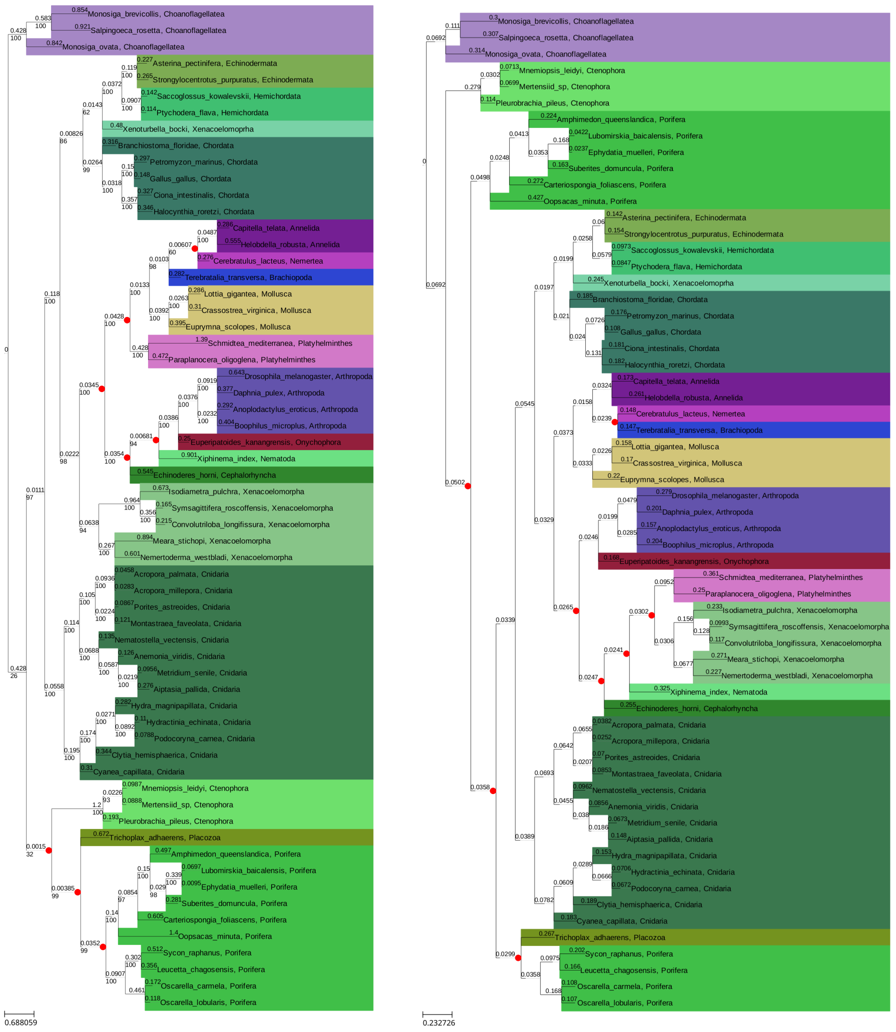

(a) ML tree of Step 3 (b) ML tree of Step 1

Figure S6: **Detailed trees for the Ryan et al. dataset.** Maximum likelihood topologies inferred by the site-heterogeneous GTR+CAT-PMSF model (a), and the site-homogeneous LG model (b) of the Ryan et al. (2013) dataset representing 60 species. Color codes are the same for species of the same taxa. Red dots mark differences between topologies (a) and (b) and correspond to the Robinson-Foulds metric (Robinson and Foulds, 1981). Values above branches represent inferred branch lengths, values below branches represent bootstrap support.

#### S3 Model performance

In this section, we present extended results with more models for all analyses performed in this work including a new way of visualizing the support between two competing topologies over the spectrum of sites with different $K_{\text{eff}}$. In the following, we briefly describe additional methods necessary to create these figures.

*$K_{eff}$ cumulative log-likelihood difference figures.* – Figures S8, S9, and S10 show cumulative log-likelihood differences. The cumulative log-likelihood difference at a specific $K_{\text{eff}}$ value is the sum of log-likelihood differences of sites of the specific $K_{\text{eff}}$ value or lower. These figures are ideal for showing the preferred topology and for determining how decisive different $K_{\text{eff}}$-ranges are. Corresponding with Section Materials and Methods, a total log-likelihood difference above zero (blue area) means support for topology $B$ and a total log-likelihood difference below zero (yellowish area) means support for the topology $A$.

*Site-proportional $K_{eff}$-axis.* – A modified version of the representation above can be seen in Figures S11 and S12 when the length of the x-axis is proportional to the number of sites observing the given $K_{\text{eff}}$ value range.

*Different bin sizes.* – Figures S13, S14 and S15 show how the compositional constraint analysis changes for different bin sizes.

*Performance of CAT-PMSF compared to the PMSF method.* – The quality of tree inference using amino acid site-specific stationary distributions estimated by the CAT-PMSF and PMSF methods were explored (Fig. S7, S8, S9, S10 and S16). The exchangeabilities and discrete rate heterogeneity category numbers matched the ones used in previous steps (e.g. LG+PMSF was used for PhyloBayes LG+CAT).

*Settings for IQ-TREE 2 required to compare site-specific log-likelihoods of different topologies.* – IQ-TREE 2 was set to output site-specific log-likelihood values (`-wslr`) and perform ultra-fast bootstrap (`-B 1000`). The output of site-specific log-likelihood is necessary in order to create the "spectral analyses" but is unnecessary if the sole aim is to infer a tree less prone to LBA.

##### *S3.1 Simulation study*

We simulated amino acid alignments with 10 000 sites exhibiting across-site compositional heterogeneity (Schrempf et al., 2020) along Felsenstein-type quartet trees (insets in top row of Figure 2; Felsenstein, 1978) with different branch lengths ($q = 0.1$ kept constant; $p = 0.1$ to $p = 2.0$ changed) under a site-heterogeneous model with varying stationary distributions and constant Poisson (Felsenstein, 1973; Nei, 1987) exchangeabilities. We performed analyses with Poisson+CAT-PMSF, LG+CAT-PMSF, GTR+CAT-PMSF, LG+C60 (Quang et al., 2008), LG+PMSF+C60 (Quang et al., 2008; Wang et al., 2018), LG (Le and Gascuel, 2008), and GTR (Tavaré, 1986) models constrained to the correct topology as well as the incorrect (Farris-type) topology (inset in bottom row of Figure 2; Farris, 1999) with IQ-TREE 2 (Minh et al., 2020). For Poisson+CAT-PMSF the effect of the guide topology used in Step 2 was explored, either setting the topology to the one inferred in Step 1 or to the genuine one. A third run uses the genuine topology and site-specific amino acid frequencies (*Poisson orig freq*). The site-specific log-likelihood differences $\Delta \log L$ between the maximum likelihood trees of the two competing topologies binned according to their effective number of amino acids $K_{\text{eff}}$ are show in Figure S7. The cumulative site-specific log-likelihood differences $\Delta \log L$ between the maximum likelihood trees of the two competing topologies as a function of the sites with increasing effective number of amino acids $K_{\text{eff}}$ estimated by PhyloBayes (Lartillot and Philippe, 2004) are shown in Figure S8. The LG, and GTR models incorrectly infer Farris-type trees if $p \geq 0.8$. The exact log-likelihood differences of the competing topologies are shown in Table S1 for Poisson+CAT-PMSF and in Table S2 for the site-homogeneous LG model.

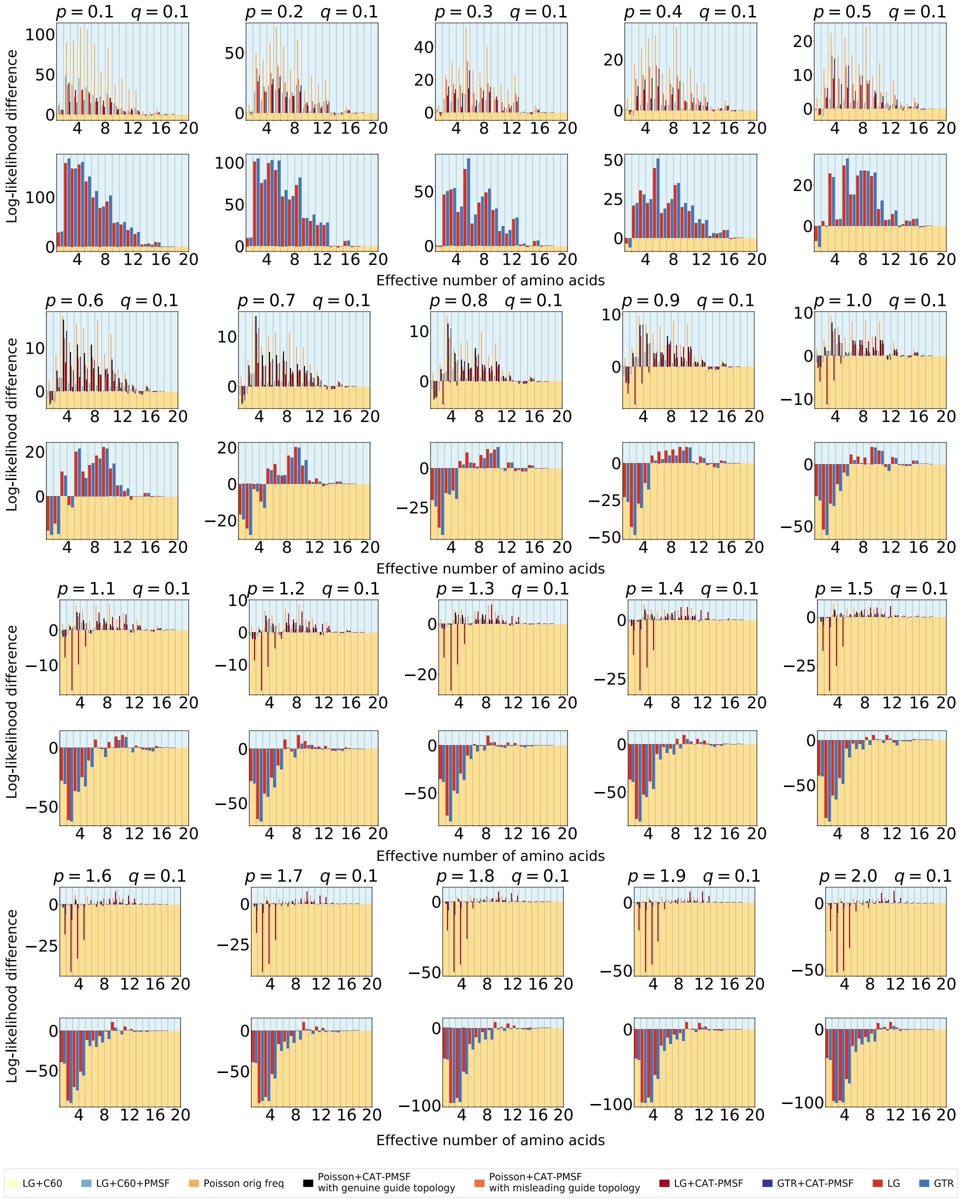

Figure S7: **Compositional constraint analysis of the simulation study, binned representation.** Extended simulation study for all branch length combinations of $p$ and $q$. The LG and GTR models incorrectly infer Farris-type trees if $p \geq 0.8$.

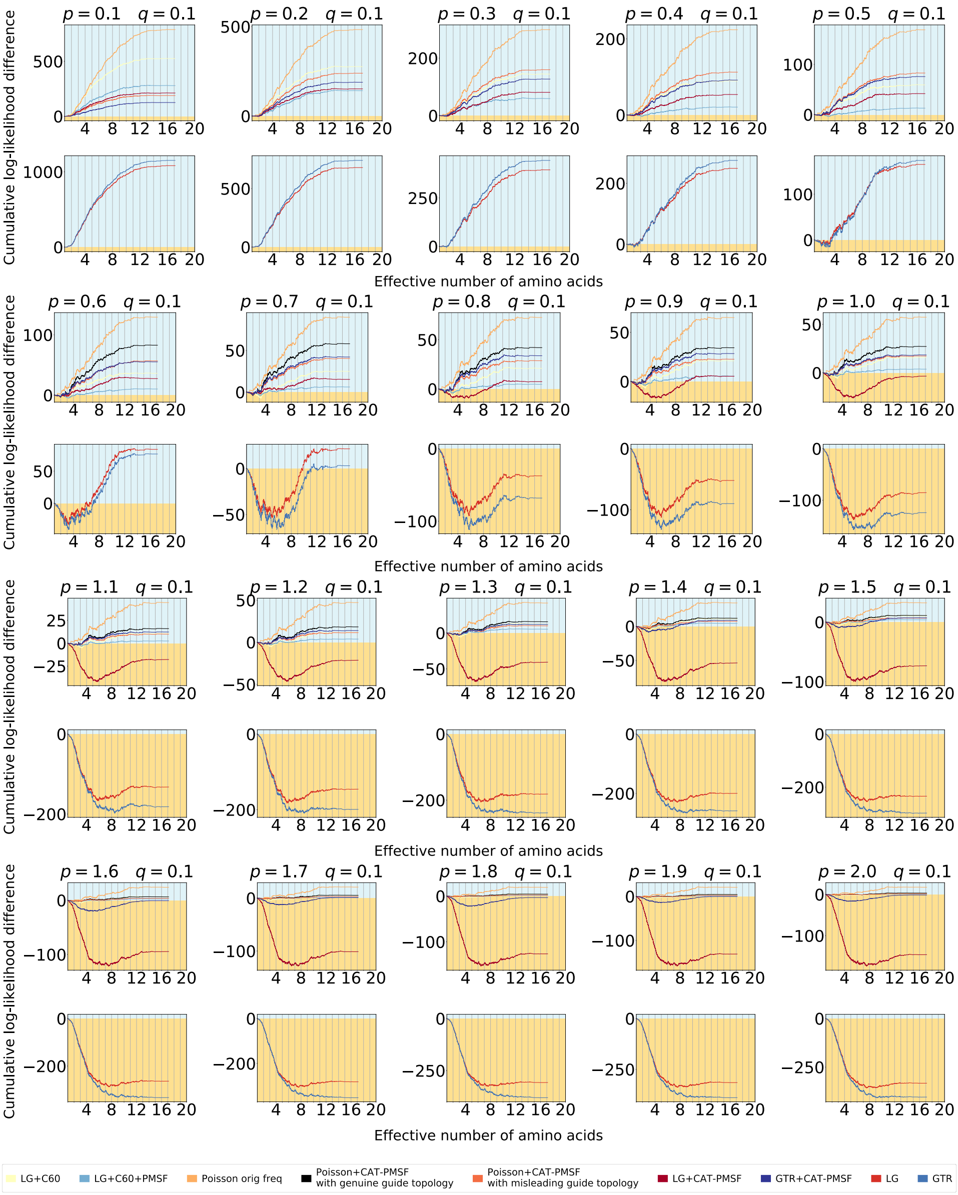

Figure S8: **Compositional constraint analysis of the simulation study, cumulative representation.** Cumulative log-likelihood differences for all branch length combinations of $p$ and $q$. The LG and GTR models incorrectly infer Farris-type trees if $p \geq 0.8$.

Table S1: **Log-likelihood results of the simulation study, Poisson+CAT-PMSF model.**

| q | p | $\log L_{\text{Farris}}$ | $\log L_{\text{Felsenstein}}$ | $\Delta \log L$ |
|---|---|---|---|---|
| 0.1 | 0.1 | -23156.9 | -22968.5 | 188.6 |
| 0.1 | 0.2 | -33954.0 | -33714.9 | 239.1 |
| 0.1 | 0.3 | -39383.5 | -39224.0 | 159.5 |
| 0.1 | 0.4 | -42175.9 | -42063.4 | 112.5 |
| 0.1 | 0.5 | -44926.5 | -44843.7 | 82.8 |
| 0.1 | 0.6 | -46463.8 | -46407.0 | 56.8 |
| 0.1 | 0.7 | -48234.8 | -48194.4 | 40.4 |
| 0.1 | 0.8 | -49813.8 | -49785.1 | 28.7 |
| 0.1 | 0.9 | -50532.3 | -50509.6 | 22.7 |
| 0.1 | 1.0 | -51238.1 | -51220.8 | 17.3 |
| 0.1 | 1.1 | -52096.2 | -52086.3 | 9.9 |
| 0.1 | 1.2 | -52876.2 | -52864.8 | 11.4 |
| 0.1 | 1.3 | -53682.7 | -53672.8 | 9.9 |
| 0.1 | 1.4 | -54222.2 | -54215.1 | 7.1 |
| 0.1 | 1.5 | -54866.1 | -54859.5 | 6.6 |
| 0.1 | 1.6 | -54976.4 | -54972.9 | 3.5 |
| 0.1 | 1.7 | -55344.2 | -55341.5 | 2.7 |
| 0.1 | 1.8 | -55692.6 | -55690.6 | 2.0 |
| 0.1 | 1.9 | -55763.7 | -55761.7 | 2.0 |
| 0.1 | 2.0 | -56078.7 | -56077.3 | 1.4 |

We simulated amino acid alignments with 10 000 sites exhibiting across-site compositional heterogeneity (Schrempf et al., 2020) along Felsenstein-type trees (insets in top row of Figure 2; Felsenstein, 1978) with different branch lengths ($q = 0.1$; $0.1 < p2.0$). Maximum log-likelihoods and log-likelihood differences of the competing topologies of the simulation dataset using the Poisson+CAT-PMSF model are shown. The true branch lengths $p$ and $q$ are indicated. A positive log-likelihood difference indicates support for the correct topology (Felsenstein-type), a negative difference for the incorrect (Farris-type) topology.

Table S2: **Log-likelihood results of the simulation study, LG model.**

| q | p | $\log L_{\text{Farris}}$ | $\log L_{\text{Felsenstein}}$ | $\Delta \log L$ |
|---|---|---|---|---|
| 0.1 | 0.1 | -53706.3 | -52625.0 | 1081.3 |
| 0.1 | 0.2 | -58628.7 | -57949.5 | 679.2 |
| 0.1 | 0.3 | -62163.4 | -61769.0 | 394.4 |
| 0.1 | 0.4 | -64806.2 | -64557.0 | 249.2 |
| 0.1 | 0.5 | -66744.9 | -66579.6 | 165.3 |
| 0.1 | 0.6 | -68552.4 | -68468.5 | 83.9 |
| 0.1 | 0.7 | -70075.2 | -70053.8 | 21.4 |
| 0.1 | 0.8 | -71390.2 | -71427.6 | -37.4 |
| 0.1 | 0.9 | -72440.1 | -72492.2 | -52.1 |
| 0.1 | 1.0 | -73454.8 | -73540.1 | -85.3 |
| 0.1 | 1.1 | -74334.2 | -74466.6 | -132.4 |
| 0.1 | 1.2 | -75057.2 | -75202.9 | -145.7 |
| 0.1 | 1.3 | -75683.1 | -75864.7 | -181.6 |
| 0.1 | 1.4 | -76326.1 | -76525.7 | -199.6 |
| 0.1 | 1.5 | -76910.3 | -77144.3 | -234.0 |
| 0.1 | 1.6 | -77480.0 | -77743.8 | -263.8 |
| 0.1 | 1.7 | -77879.9 | -78161.7 | -281.8 |
| 0.1 | 1.8 | -78269.3 | -78576.8 | -307.5 |
| 0.1 | 1.9 | -78647.1 | -78962.5 | -315.4 |
| 0.1 | 2.0 | -79045.8 | -79378.8 | -333.0 |

We simulated amino acid alignments with 10 000 sites exhibiting across-site compositional heterogeneity (Schrempf et al., 2020) along Felsenstein-type trees (insets in top row of Figure 2; Felsenstein, 1978) with different branch lengths ($q = 0.1$; $0.1 < p < 2.0$). Maximum log-likelihoods and log-likelihood differences of the competing topologies of the simulation dataset using the LG model are shown. The true branch lengths $p$ and $q$ are indicated. A positive log-likelihood difference indicates support for the correct topology (Felsenstein-type), a negative difference for the incorrect (Farris-type) topology.

##### *S3.2 Empirical datasets*

Here we show extended compositional constraint analysis figures with more models and with cumulative log-likelihoods (Fig. S9 and S10), site-proportional $K_{\mathrm{eff}}$ (Fig. S11 and S12) or different bin sizes (Fig. S13, S14 and S15).

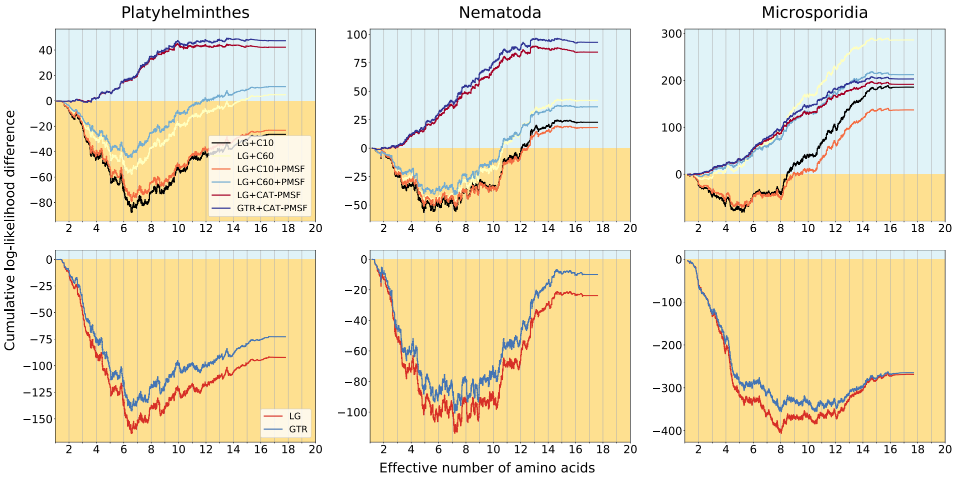

Figure S9: **Compositional constraint analysis of the empirical datasets, cumulative representation.** We analyzed three empirical datasets including (left) Platyhelminthes and (middle) Nematoda (Philippe et al., 2005), and (right) Microsporidia (Brinkmann et al., 2005). We performed analyses with LG+CAT-PMSF, GTR+CAT-PMSF, LG+C10 and LG+C60 (Quang et al., 2008), LG+C10+PMSF and LG+C60+PMSF (Quang et al., 2008; Wang et al., 2018), LG (Le and Gascuel, 2008) and GTR (Tavaré, 1986) models constrained to either one of two competing topologies with IQ-TREE 2 (Minh et al., 2020). The blue background indicates support for the topologies in Figures S3a, S1a and S2a; the yellowish background for the long branch attraction-prone topologies in Figures S3b, S1b and S2b. The cumulative log-likelihood differences $\Delta \log L$ between the LBA-prone and the non-LBA-prone topologies as a function of the sites with increasing effective number of amino acids $K_{\mathrm{eff}}$ estimated by PhyloBayes (Lartillot and Philippe, 2004) are shown. Site-homogeneous models infer the topology prone to long branch attraction for all three datasets. Conventional site-heterogeneous models infer the topology purportedly free of long branch attraction when enough categories are used. For the Platyhelminthes dataset 10 categories are not enough, but 60 are. For the Nematoda and Microsporidia datasets 10 categories are sufficient to model the across-site compositional heterogeneity in the data. We observe bias for the long branch attraction-prone topology for the low $K_{\mathrm{eff}}$ sites when using these methods, except for Microsporidia when using 60 categories. The CAT-PMSF method on the other hand is not showing correlations between $K_{\mathrm{eff}}$ value and topology preference.

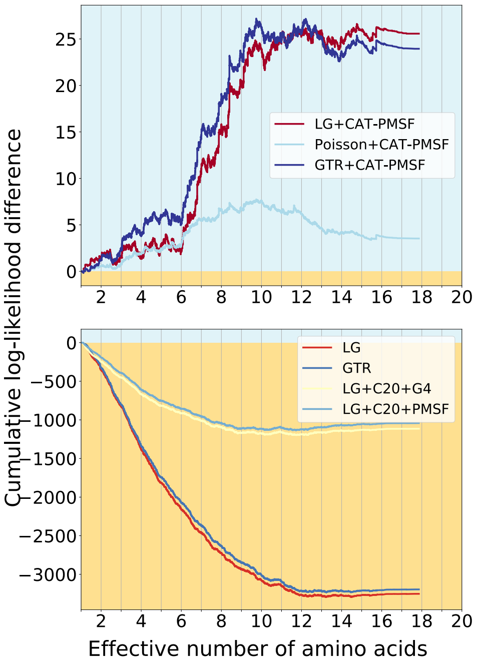

Figure S10: **Compositional constraint analysis of Simion et al.'s reduced outgroup dataset, cumulative representation.** We performed analyses with Poisson+CAT-PMSF, LG+CAT-PMSF, GTR+CAT-PMSF, LG (Le and Gascuel, 2008), GTR (Tavaré, 1986), LG+C20+PMSF (Quang et al., 2008; Wang et al., 2018) and LG+C20 (Quang et al., 2008) models constrained to either one of two competing topologies (the blue background supports the topology in Figure S4a, the yellowish background the topology in Figure S4b) with IQ-TREE 2 (Minh et al., 2020). The cumulative site-specific log-likelihood differences $\Delta \log L$ between the maximum likelihood trees of the two competing topologies as a function of the sites with increasing effective number of amino acids $K_{\text{eff}}$ estimated by PhyloBayes (Lartillot and Philippe, 2004) are shown. Site-homogeneous models and the site-heterogeneous LG+C20+PMSF and LG+C20 models show inconsistent signal between constrained versus unconstrained sites and favor Ctenophora at the animal root. CAT-PMSF favors Porifera at the animal root although this result is only significant when using the closest outgroup exclusively.

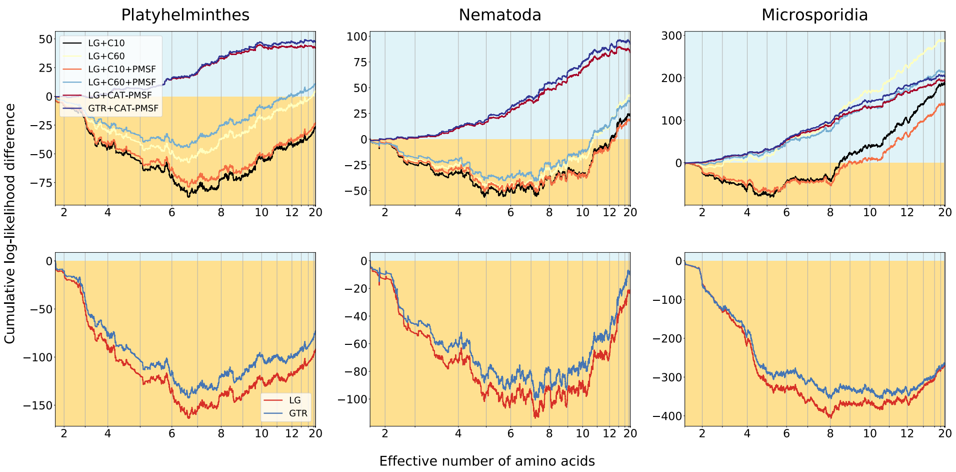

Figure S11: **Compositional constraint analysis of the empirical datasets, cumulative representation, with site-proportional $K_{\mathbf{eff}}$.** Site-proportional version of S9 to visualise the number of sites within a given $K_{\text{eff}}$ range.

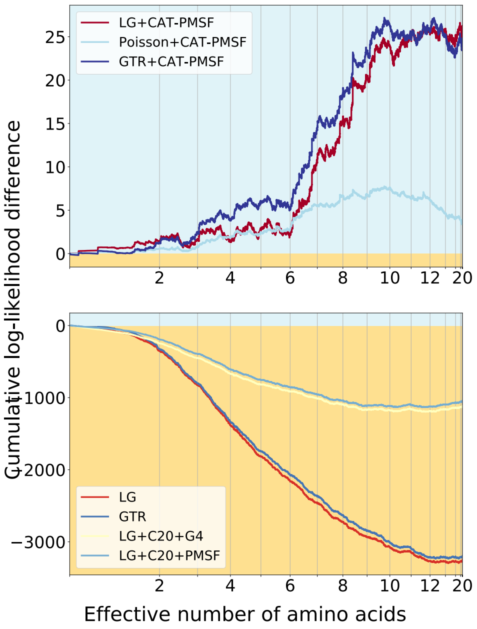

Figure S12: **Compositional constraint analysis of Simion et al.'s reduced outgroup dataset, cumulative representation, with site-proportional $K_{\mathbf{eff}}$.** Site-proportional version of S10 to visualise the number of sites within a given $K_{\text{eff}}$ range.

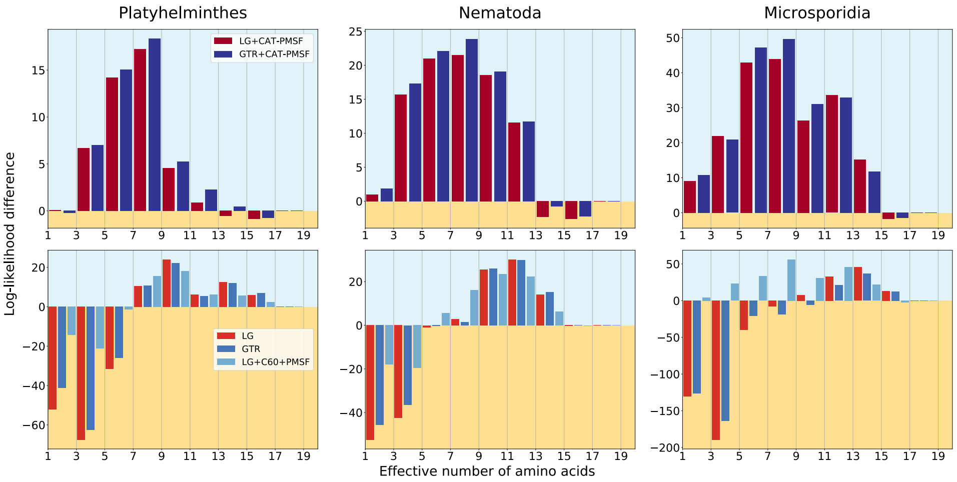

Figure S13: **Variation of compositional constraint analysis of the empirical datasets on Figure 3 with 10 bins.**

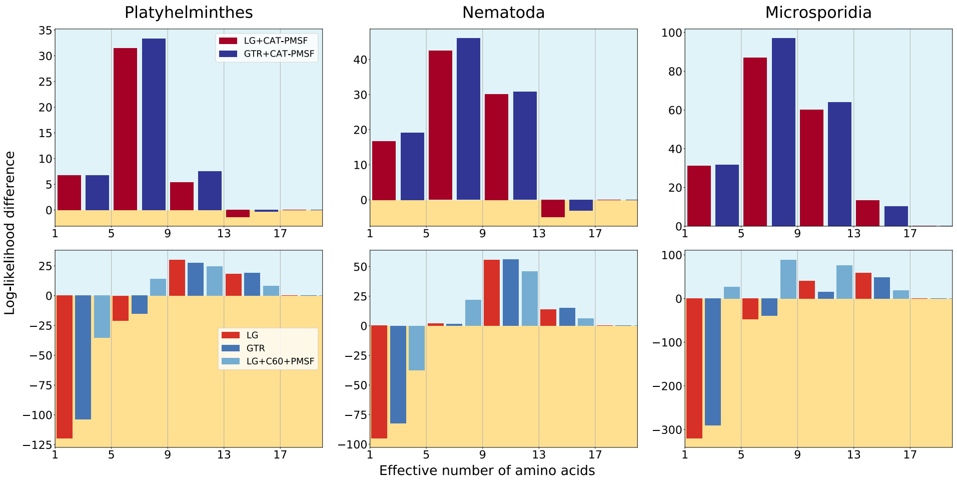

Figure S14: **Variation of compositional constraint analysis of the empirical datasets on Figure 3 with 5 bins.**

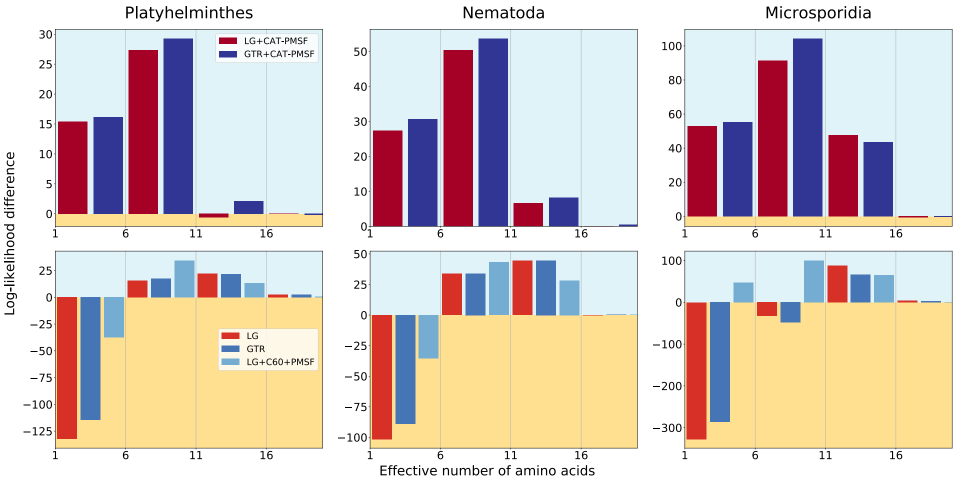

Figure S15: **Variation of compositional constraint analysis of the empirical datasets on Figure 3 with 4 bins.**

#### S4 Measuring correlation between $K_{\text{EFF}}$ and site-specific log-likelihood difference

To assess the overall significance of the site-specific log-likelihood differences and to evaluate whether site-homogeneous models are indeed biased, we calculated the correlation between the log-likelihood difference and the effective number of used amino acids $K_{\text{eff}}$ per site. To this aim, the Pearson's correlation coefficient and p-value were calculated for all datasets and models (Tables S3 and S4). Further, the rank correlation coefficients of two nonparametric tests, the Spearman's rank (Tables S6 and S5) and the Kendall's rank (Tables S8 and S7) were also calculated usign SciPy's libraries (Jones et al., 2001).

Overall, we observe: Invariant sites or sites with $K_{\text{eff}}$ below 3 have low site-specific log-likelihood differences especially for analyses based on CAT-PMSF. Thereby, the rank correlation coefficients tend to be significant. If we remove sites with $K_{\text{eff}}$ below 3, the rank correlation coefficients are not anymore significant for analyses based on CAT-PMSF (data not shown). Some sites with $3 < K_{\text{eff}} < 7$ show high site-specific log-likelihood differences with classical models. They greatly influence the topology preference but not so much the rank correlation coefficients. For these reasons, we think that correlation coefficients based on the site-specific log-likelihood differences such as the Pearson's coefficient should be preferred over correlation coefficients solely based on the rank.

##### *S4.1 Pearson's correlation coefficients and p-values*

Table S3: **Pearson's correlation test restults for the simulation study.**

| Dataset | Model | Pearson's $r$ | $p$-value |
|---|---|---|---|
| $\mathbf{p = 0.2, q = 0.1}$ | **Poisson+CAT-PMSF** | **0.021** | **0.032** |
| $p = 0.2, q = 0.1$ | LG+CAT-PMSF | 0.012 | 0.246 |
| $p = 0.2, q = 0.1$ | GTR+CAT-PMSF | 0.016 | 0.121 |
| $p = 0.2, q = 0.1$ | LG+PMSF+C60 | 0.018 | 0.079 |
| $\mathbf{p = 0.2, q = 0.1}$ | **LG** | **0.026** | **0.009** |
| $\mathbf{p = 0.2, q = 0.1}$ | **GTR** | **0.033** | $\mathbf{9.553 \cdot 10^{-4}}$ |
| $p = 0.8, q = 0.1$ | Poisson+CAT-PMSF | 0.013 | 0.179 |
| $\mathbf{p = 0.8, q = 0.1}$ | **LG+CAT-PMSF** | **0.043** | $\mathbf{1.565 \cdot 10^{-5}}$ |
| $p = 0.8, q = 0.1$ | GTR+CAT-PMSF | 0.011 | 0.276 |
| $p = 0.8, q = 0.1$ | LG+PMSF+C60 | 0.016 | 0.107 |
| $\mathbf{p = 0.8, q = 0.1}$ | **LG** | **0.051** | $\mathbf{3.520 \cdot 10^{-7}}$ |
| $\mathbf{p = 0.8, q = 0.1}$ | **GTR** | **0.045** | $\mathbf{7.169 \cdot 10^{-6}}$ |
| $p = 1.2, q = 0.1$ | Poisson+CAT-PMSF | 0.013 | 0.207 |
| $\mathbf{p = 1.2, q = 0.1}$ | **LG+CAT-PMSF** | **0.096** | $\mathbf{9.348 \cdot 10^{-22}}$ |
| $p = 1.2, q = 0.1$ | GTR+CAT-PMSF | 0.012 | 0.236 |
| $p = 1.2, q = 0.1$ | LG+PMSF+C60 | 0.017 | 0.084 |
| $\mathbf{p = 1.2, q = 0.1}$ | **LG** | **0.072** | $\mathbf{4.325 \cdot 10^{-13}}$ |
| $\mathbf{p = 1.2, q = 0.1}$ | **GTR** | **0.053** | $\mathbf{1.093 \cdot 10^{-7}}$ |

Pearson's $r$ and associated $p$-values for the correlation between the effective number of used amino acids $K_{\text{eff}}$ per site and the site-specific log-likelihood difference between the two competing topologies. Values for the simulation study and models described in the main text are shown. Bold rows show significant correlation.

Table S4: **Pearson's correlation test restults for the empirical datasets.**

| Dataset | Model | Pearson's $r$ | $p$-value |
|---|---|---|---|
| Platyhelminthes | LG+CAT-PMSF | $-0.003$ | 0.640 |
| Platyhelminthes | GTR+CAT-PMSF | $1.135 \cdot 10^{-5}$ | 0.998 |
| **Platyhelminthes** | **LG+PMSF+C60** | **0.024** | $\mathbf{7.241 \cdot 10^{-6}}$ |
| **Platyhelminthes** | **LG** | **0.028** | $\mathbf{1.093 \cdot 10^{-7}}$ |
| **Platyhelminthes** | **GTR** | **0.026** | $\mathbf{1.015 \cdot 10^{-6}}$ |
| Nematoda | LG+CAT-PMSF | 0.008 | 0.118 |
| Nematoda | GTR+CAT-PMSF | 0.009 | 0.077 |
| **Nematoda** | **LG+PMSF+C60** | **0.025** | $\mathbf{2.504 \cdot 10^{-6}}$ |
| **Nematoda** | **LG** | **0.023** | $\mathbf{1.234 \cdot 10^{-5}}$ |
| **Nematoda** | **GTR** | **0.022** | $\mathbf{2.514 \cdot 10^{-5}}$ |
| Microsporidia | LG+CAT-PMSF | 0.011 | 0.099 |
| Microsporidia | GTR+CAT-PMSF | 0.009 | 0.147 |
| **Microsporidia** | **LG+PMSF+C60** | **0.018** | **0.005** |
| **Microsporidia** | **LG** | **0.044** | $\mathbf{5.931 \cdot 10^{-12}}$ |
| **Microsporidia** | **GTR** | **0.040** | $\mathbf{4.244 \cdot 10^{-10}}$ |
| Simion reduced outgroup | LG+CAT-PMSF | 0.002 | 0.201 |
| Simion reduced outgroup | GTR+CAT-PMSF | $3.079 \cdot 10^{-4}$ | 0.845 |
| **Simion reduced outgroup** | **LG+PMSF+C20** | **0.014** | $\mathbf{9.432 \cdot 10^{-20}}$ |
| **Simion reduced outgroup** | **LG** | **0.008** | $\mathbf{2.836 \cdot 10^{-7}}$ |
| **Simion reduced outgroup** | **GTR** | **0.007** | $\mathbf{8.318 \cdot 10^{-6}}$ |
| **Ryan** | **CAT-PMSF LG** | $\mathbf{-0.015}$ | $\mathbf{1.024 \cdot 10^{-5}}$ |
| **Ryan** | **CAT-PMSF GTR** | $\mathbf{-0.013}$ | $\mathbf{1.696 \cdot 10^{-4}}$ |
| **Ryan** | **LG+PMSF+C60** | **0.011** | $\mathbf{8.659 \cdot 10^{-4}}$ |
| **Ryan** | **LG** | **0.011** | $\mathbf{1.070 \cdot 10^{-3}}$ |
| **Ryan** | **GTR** | **0.010** | $\mathbf{4.278 \cdot 10^{-3}}$ |

Pearson's $r$ and associated $p$-values for the correlation between the site-specific effective number of used amino acids $K_{\text{eff}}$ as estimated by PhyloBayes (Lartillot and Philippe, 2004) and the site-specific log-likelihood difference between the two competing topologies. Results for the empirical datasets and models are shown. Bold rows show significant correlation.

##### *S4.2 Spearman's rank correlation coefficients and p-values*

Table S5: **Spearman's rank correlation test results for the simulation.**

| Dataset | Model | Spearman's $\rho$ | $p$-value |
|---|---|---|---|
| $\mathbf{p = 0.2, q = 0.1}$ | **Poisson+CAT-PMSF** | $\mathbf{-0.005}$ | **0.602** |
| $p = 0.2, q = 0.1$ | LG+CAT-PMSF | $-0.078$ | $4.754 \cdot 10^{-15}$ |
| $p = 0.2, q = 0.1$ | GTR+CAT-PMSF | $-0.038$ | $1.220 \cdot 10^{-4}$ |
| $p = 0.2, q = 0.1$ | LG+PMSF+C60 | $-0.040$ | $6.921 \cdot 10^{-5}$ |
| $p = 0.2, q = 0.1$ | LG | $-0.048$ | $1.504 \cdot 10^{-6}$ |
| $\mathbf{p = 0.2, q = 0.1}$ | **GTR** | $\mathbf{-0.018}$ | **0.073** |
| $p = 0.8, q = 0.1$ | Poisson+CAT-PMSF | $-0.045$ | $5.996 \cdot 10^{-6}$ |
| $p = 0.8, q = 0.1$ | LG+CAT-PMSF | 0.075 | $6.967 \cdot 10^{-14}$ |
| $p = 0.8, q = 0.1$ | GTR+CAT-PMSF | $-0.046$ | $4.476 \cdot 10^{-6}$ |
| $\mathbf{p = 0.8, q = 0.1}$ | **LG+PMSF+C60** | $\mathbf{-0.010}$ | **0.304** |
| $p = 0.8, q = 0.1$ | LG | 0.115 | $1.605 \cdot 10^{-30}$ |
| $p = 0.8, q = 0.1$ | GTR | 0.114 | $4.547 \cdot 10^{-30}$ |
| $p = 1.2, q = 0.1$ | Poisson+CAT-PMSF | $-0.038$ | $1.597 \cdot 10^{-4}$ |
| $p = 1.2, q = 0.1$ | LG+CAT-PMSF | 0.164 | $1.531 \cdot 10^{-61}$ |
| $p = 1.2, q = 0.1$ | GTR+CAT-PMSF | $-0.045$ | $6.292 \cdot 10^{-6}$ |
| $\mathbf{p = 1.2, q = 0.1}$ | **LG+PMSF+C60** | $\mathbf{-0.013}$ | **0.205** |
| $p = 1.2, q = 0.1$ | LG | 0.147 | $3.294 \cdot 10^{-49}$ |
| $p = 1.2, q = 0.1$ | GTR | 0.138 | $1.931 \cdot 10^{-43}$ |

Bold rows show non-significant correlations.

Table S6: **Spearman's rank correlation test results for the empirical datasets.**

| Dataset | Model | Spearman's $\rho$ | $p$-value |
|---|---|---|---|
| Platyhelminthes | LG+CAT-PMSF | 0.016 | 0.002 |
| **Platyhelminthes** | **GTR+CAT-PMSF** | **0.005** | **0.327** |
| Platyhelminthes | LG+PMSF+C60 | 0.043 | $3.605 \cdot 10^{-16}$ |
| Platyhelminthes | LG | 0.097 | $1.654 \cdot 10^{-74}$ |
| Platyhelminthes | GTR | 0.089 | $3.132 \cdot 10^{-63}$ |
| Nematoda | LG+CAT-PMSF | 0.035 | $3.703 \cdot 10^{-11}$ |
| Nematoda | GTR+CAT-PMSF | 0.025 | $1.864 \cdot 10^{-6}$ |
| Nematoda | LG+PMSF+C60 | 0.026 | $7.573 \cdot 10^{-7}$ |
| Nematoda | LG | 0.028 | $1.086 \cdot 10^{-7}$ |
| Nematoda | GTR | 0.029 | $7.375 \cdot 10^{-8}$ |
| Microsporidia | LG+CAT-PMSF | 0.038 | $2.261 \cdot 10^{-9}$ |
| Microsporidia | GTR+CAT-PMSF | 0.017 | 0.008 |
| Microsporidia | LG+PMSF+C60 | 0.038 | $2.452 \cdot 10^{-9}$ |
| Microsporidia | LG | 0.144 | $2.725 \cdot 10^{-113}$ |
| Microsporidia | GTR | 0.138 | $5.984 \cdot 10^{-104}$ |
| Simion reduced outgroup | LG+CAT-PMSF | 0.013 | $2.802 \cdot 10^{-16}$ |
| Simion reduced outgroup | GTR+CAT-PMSF | 0.072 | 0.0 |
| Simion reduced outgroup | LG+PMSF+C20 | 0.006 | $3.481 \cdot 10^{-4}$ |
| Simion reduced outgroup | LG | 0.008 | $9.191 \cdot 10^{-8}$ |
| Simion reduced outgroup | GTR | 0.043 | $7.189 \cdot 10^{-164}$ |
| Ryan | LG+CAT-PMSF | −0.116 | $2.180 \cdot 10^{-260}$ |
| Ryan | GTR+CAT-PMSF | −0.068 | $9.809 \cdot 10^{-90}$ |
| Ryan | LG+PMSF+C60 | 0.115 | $1.352 \cdot 10^{-255}$ |
| Ryan | LG | 0.111 | $1.763 \cdot 10^{-239}$ |
| Ryan | GTR | 0.105 | $1.414 \cdot 10^{-213}$ |

Bold rows show non-significant correlation.

##### *S4.3 Kendall's rank correlation coefficients and p-values*

Table S7: **Kendall's rank correlation test results for the simulation.**

| Dataset | Model | Kendall's $\tau$ | $p$-value |
|---|---|---|---|
| $\mathbf{p = 0.2, q = 0.1}$ | **Poisson+CAT-PMSF** | $\mathbf{-0.004}$ | $\mathbf{0.600}$ |
| $p = 0.2, q = 0.1$ | LG+CAT-PMSF | $-0.053$ | $3.964 \cdot 10^{-15}$ |
| $p = 0.2, q = 0.1$ | GTR+CAT-PMSF | $-0.026$ | $1.045 \cdot 10^{-4}$ |
| $p = 0.2, q = 0.1$ | LG+PMSF+C60 | $-0.027$ | $6.627 \cdot 10^{-5}$ |
| $p = 0.2, q = 0.1$ | LG | $-0.032$ | $1.665 \cdot 10^{-6}$ |
| $\mathbf{p = 0.2, q = 0.1}$ | **GTR** | $\mathbf{-0.012}$ | $\mathbf{0.071}$ |
| $p = 0.8, q = 0.1$ | Poisson+CAT-PMSF | $-0.031$ | $4.956 \cdot 10^{-6}$ |
| $p = 0.8, q = 0.1$ | LG+CAT-PMSF | $0.051$ | $1.934 \cdot 10^{-14}$ |
| $p = 0.8, q = 0.1$ | GTR+CAT-PMSF | $-0.031$ | $4.828 \cdot 10^{-6}$ |
| $\mathbf{p = 0.8, q = 0.1}$ | **LG+PMSF+C60** | $\mathbf{-0.007}$ | $\mathbf{0.281}$ |
| $p = 0.8, q = 0.1$ | LG | $0.078$ | $6.343 \cdot 10^{-31}$ |
| $p = 0.8, q = 0.1$ | GTR | $0.077$ | $1.104 \cdot 10^{-30}$ |
| $p = 1.2, q = 0.1$ | Poisson+CAT-PMSF | $-0.026$ | $9.349 \cdot 10^{-5}$ |
| $p = 1.2, q = 0.1$ | LG+CAT-PMSF | $0.112$ | $9.416 \cdot 10^{-63}$ |
| $p = 1.2, q = 0.1$ | GTR+CAT-PMSF | $-0.031$ | $4.741 \cdot 10^{-6}$ |
| $\mathbf{p = 1.2, q = 0.1}$ | **LG+PMSF+C60** | $\mathbf{-0.009}$ | $\mathbf{0.183}$ |
| $p = 1.2, q = 0.1$ | LG | $0.099$ | $1.062 \cdot 10^{-49}$ |
| $p = 1.2, q = 0.1$ | GTR | $0.093$ | $3.651 \cdot 10^{-44}$ |

Bold rows show non-significant correlation.

Table S8: **Kendall's rank correlation test results for the empirical datasets.**

| Dataset | Model | Kendall's $\tau$ | $p$-value |
|---|---|---|---|
| Platyhelminthes | LG+CAT-PMSF | 0.019 | $9.877 \cdot 10^{-8}$ |
| Platyhelminthes | GTR+CAT-PMSF | 0.010 | 0.004 |
| Platyhelminthes | LG+PMSF+C60 | 0.027 | $1.183 \cdot 10^{-13}$ |
| Platyhelminthes | LG | 0.064 | $6.404 \cdot 10^{-73}$ |
| Platyhelminthes | GTR | 0.061 | $1.148 \cdot 10^{-66}$ |
| Nematoda | LG+CAT-PMSF | 0.047 | $3.472 \cdot 10^{-40}$ |
| Nematoda | GTR+CAT-PMSF | 0.039 | $2.030 \cdot 10^{-28}$ |
| **Nematoda** | **LG+PMSF+C60** | **0.001** | **0.687** |
| **Nematoda** | **LG** | **0.005** | **0.184** |
| **Nematoda** | **GTR** | $\mathbf{1.770 \cdot 10^{-4}}$ | **0.960** |
| Microsporidia | LG+CAT-PMSF | 0.035 | $4.865 \cdot 10^{-16}$ |
| Microsporidia | GTR+CAT-PMSF | 0.018 | $2.427 \cdot 10^{-5}$ |
| Microsporidia | LG+PMSF+C60 | 0.024 | $1.592 \cdot 10^{-8}$ |
| Microsporidia | LG | 0.090 | $3.567 \cdot 10^{-98}$ |
| Microsporidia | GTR | 0.087 | $8.006 \cdot 10^{-91}$ |
| Simion reduced outgroup | LG+CAT-PMSF | 0.018 | $2.376 \cdot 10^{-60}$ |
| Simion reduced outgroup | GTR+CAT-PMSF | 0.074 | 0.0 |
| Simion reduced outgroup | LG+PMSF+C20 | −0.028 | $1.118 \cdot 10^{-157}$ |
| Simion reduced outgroup | LG | −0.029 | $2.112 \cdot 10^{-165}$ |
| Simion reduced outgroup | GTR | 0.009 | $2.045 \cdot 10^{-16}$ |
| Ryan | LG+CAT-PMSF | −0.080 | $4.908 \cdot 10^{-256}$ |
| Ryan | GTR+CAT-PMSF | −0.047 | $3.986 \cdot 10^{-83}$ |
| Ryan | LG+PMSF+C60 | 0.064 | $2.730 \cdot 10^{-176}$ |
| Ryan | LG | 0.053 | $2.898 \cdot 10^{-124}$ |
| Ryan | GTR | 0.049 | $1.198 \cdot 10^{-106}$ |

Bold rows show non-significant correlation.

#### S5 AU TESTS

Approximately Unbiased tests (AU, Shimodaira, 2002) were run on the maximum likelihood trees constrained to the topologies inferred at Step 1 and Step 3 of the CAT-PMSF method. As shown in Tables S12, S13, S14 and S15 the topology inferred at Step 3 (T1) is significantly more probable under the model than the LBA-prone topology of Step 1 (T2). This is in accordance with the results presented by the compositional constraint analysis.

Table S9: **AU test results for the simulation study with GTR+CAT-PMSF model.**

| q | p | p-$AU_{T1}$ | | p-$AU_{T2}$ | |
|---|---|---|---|---|---|
| 0.1 | 0.1 | 1 | + | $1.87 \cdot 10^{-8}$ | − |
| 0.1 | 0.2 | 1 | + | $4.8 \cdot 10^{-58}$ | − |
| 0.1 | 0.3 | 1 | + | $6.75 \cdot 10^{-5}$ | − |
| 0.1 | 0.4 | 1 | + | $1.17 \cdot 10^{-49}$ | − |
| 0.1 | 0.5 | 1 | + | $2.87 \cdot 10^{-6}$ | − |
| 0.1 | 0.6 | 1 | + | $6.03 \cdot 10^{-6}$ | − |
| 0.1 | 0.7 | 1 | + | $1.89 \cdot 10^{-7}$ | − |
| 0.1 | 0.8 | 1 | + | $2.31 \cdot 10^{-4}$ | − |
| 0.1 | 0.9 | 1 | + | $1.98 \cdot 10^{-4}$ | − |
| 0.1 | 1.0 | 0.998 | + | $1.67 \cdot 10^{-3}$ | − |
| 0.1 | 1.1 | 0.993 | + | $6.60 \cdot 10^{-3}$ | − |
| 0.1 | 1.2 | 0.996 | + | $3.86 \cdot 10^{-3}$ | − |
| 0.1 | 1.3 | 0.99 | + | $9.78 \cdot 10^{-3}$ | − |
| 0.1 | 1.4 | 0.964 | + | $3.56 \cdot 10^{-2}$ | − |
| 0.1 | 1.5 | 0.918 | + | $8.17 \cdot 10^{-2}$ | + |
| 0.1 | 1.6 | 0.321 | + | 0.679 | + |
| 0.1 | 1.7 | 0.54 | + | 0.46 | + |
| 0.1 | 1.8 | 0.161 | + | 0.839 | + |
| 0.1 | 1.9 | 0.377 | + | 0.623 | + |
| 0.1 | 2.0 | 0.237 | + | 0.763 | + |

The T1 topology constraint is the same as the ML topology output by the CAT-PMSF pipeline, whereas the T2 topology constraint is the ML topology under the specified model with a constraint producing the long branch attraction artifact. Minus signs mean significant exclusion of the topology, plus signs mean that the topology could not be excluded within the 95% confidence interval.

Table S10: **AU test results for the simulation study with LG+CAT-PMSF model.**

| q | p | p-$AU_{T1}$ | | p-$AU_{T2}$ | |
|---|---|---|---|---|---|
| 0.1 | 0.1 | 1 | + | $3.06 \cdot 10^{-51}$ | − |
| 0.1 | 0.2 | 1 | + | $4.37 \cdot 10^{-48}$ | − |
| 0.1 | 0.3 | 1 | + | $3.98 \cdot 10^{-38}$ | − |
| 0.1 | 0.4 | 1 | + | $1.78 \cdot 10^{-50}$ | − |
| 0.1 | 0.5 | 1 | + | $7.64 \cdot 10^{-7}$ | − |
| 0.1 | 0.6 | 1 | + | $2.13 \cdot 10^{-4}$ | − |
| 0.1 | 0.7 | 0.998 | + | $2.34 \cdot 10^{-3}$ | − |
| 0.1 | 0.8 | 0.912 | + | $8.76 \cdot 10^{-2}$ | + |
| 0.1 | 0.9 | 0.795 | + | 0.205 | + |
| 0.1 | 1.0 | 0.255 | + | 0.745 | + |
| 0.1 | 1.1 | $3.39 \cdot 10^{-4}$ | − | 0.997 | + |
| 0.1 | 1.2 | $5.85 \cdot 10^{-4}$ | − | 0.999 | + |
| 0.1 | 1.3 | $3.87 \cdot 10^{-5}$ | − | 1 | + |
| 0.1 | 1.4 | $2.26 \cdot 10^{-6}$ | − | 1 | + |
| 0.1 | 1.5 | $4.95 \cdot 10^{-11}$ | − | 1 | + |
| 0.1 | 1.6 | $6.79 \cdot 10^{-51}$ | − | 1 | + |
| 0.1 | 1.7 | $2.11 \cdot 10^{-4}$ | − | 1 | + |
| 0.1 | 1.8 | $5.46 \cdot 10^{-4}$ | − | 0.999 | + |
| 0.1 | 1.9 | $7.35 \cdot 10^{-4}$ | − | 0.999 | + |
| 0.1 | 2.0 | $1.07 \cdot 10^{-10}$ | − | 1 | + |

The T1 topology constraint is the same as the ML topology output by the CAT-PMSF pipeline, whereas the T2 topology constraint is the ML topology under the specified model with a constraint producing the long branch attraction artifact. Minus signs mean significant exclusion of the topology, plus signs mean that the topology could not be excluded within the 95% confidence interval.

Table S11: **AU test results for the simulation study with Poisson+CAT-PMSF model.**

| q | p | p-AU$_{T1}$ | | p-AU$_{T2}$ | |
|---|---|---|---|---|---|
| 0.1 | 0.1 | 1 | + | $1.82 \cdot 10^{-8}$ | − |
| 0.1 | 0.2 | 1 | + | $2.57 \cdot 10^{-4}$ | − |
| 0.1 | 0.3 | 1 | + | $1.13 \cdot 10^{-6}$ | − |
| 0.1 | 0.4 | 1 | + | $2.59 \cdot 10^{-65}$ | − |
| 0.1 | 0.5 | 1 | + | $1.37 \cdot 10^{-62}$ | − |
| 0.1 | 0.6 | 1 | + | $2.09 \cdot 10^{-9}$ | − |
| 0.1 | 0.7 | 1 | + | $2.29 \cdot 10^{-9}$ | − |
| 0.1 | 0.8 | 1 | + | $1.46 \cdot 10^{-4}$ | − |
| 0.1 | 0.9 | 0.999 | + | $1.17 \cdot 10^{-3}$ | − |
| 0.1 | 1.0 | 0.997 | + | $3.15 \cdot 10^{-3}$ | − |
| 0.1 | 1.1 | 0.984 | + | $1.61 \cdot 10^{-2}$ | − |
| 0.1 | 1.2 | 0.989 | + | $1.07 \cdot 10^{-2}$ | − |
| 0.1 | 1.3 | 0.982 | + | $1.84 \cdot 10^{-2}$ | − |
| 0.1 | 1.4 | 0.968 | + | $3.17 \cdot 10^{-2}$ | − |
| 0.1 | 1.5 | 0.957 | + | $4.27 \cdot 10^{-2}$ | − |
| 0.1 | 1.6 | 0.898 | + | 0.102 | + |
| 0.1 | 1.7 | 0.875 | + | 0.125 | + |
| 0.1 | 1.8 | 0.835 | + | 0.165 | + |
| 0.1 | 1.9 | 0.836 | + | 0.164 | + |
| 0.1 | 2.0 | 0.786 | + | 0.214 | + |

The T1 topology constraint is the same as the ML topology output by the CAT-PMSF pipeline, whereas the T2 topology constraint is the ML topology under the specified model with a constraint producing the long branch attraction artifact. Minus signs mean significant exclusion of the topology, plus signs mean that the topology could not be excluded within the 95% confidence interval.

Table S12: **AU test results for the Microsporidia dataset with CAT-PMSF models.**

| Model | p-$AU_{T1}$ | | p-$AU_{T2}$ | |
|---|---|---|---|---|
| LG+CAT-PMSF, chain 1 | 1 | + | $1.13 \cdot 10^{-4}$ | − |
| LG+CAT-PMSF, chain 2 | 1 | + | $1.69 \cdot 10^{-7}$ | − |
| GTR+CAT-PMSF, chain 1 | 1 | + | $1.25 \cdot 10^{-49}$ | − |
| GTR+CAT-PMSF, chain 2 | 1 | + | $3.27 \cdot 10^{-74}$ | − |

The T1 topology constraint is the same as the ML topology output by the CAT-PMSF pipeline, whereas the T2 topology constraint is the ML topology under the specified model with a constraint producing the long branch attraction artifact. Minus signs mean significant exclusion of the topology, plus signs mean that the topology could not be excluded within the 95% confidence interval.

Table S13: **AU test results for the Nematoda dataset with CAT-PMSF models.**

| Model | p-$AU_{T1}$ | | p-$AU_{T2}$ | |
|---|---|---|---|---|
| LG+CAT-PMSF chain 1 | 0.484 | + | $7.18 \cdot 10^{-8}$ | − |
| LG+CAT-PMSF chain 2 | 0.51 | + | $2.02 \cdot 10^{-71}$ | − |
| GTR+CAT-PMSF chain 1 | 0.491 | + | $1.01 \cdot 10^{-10}$ | − |
| GTR+CAT-PMSF chain 2 | 0.503 | + | $1.3 \cdot 10^{-83}$ | − |

T1 tree constraint is the same topology as the ML tree outputted by the CAT-PMSF pipeline, whereas T2 tree constraint is the ML tree under the specified model with a constraint producing the long branch attraction artifact. Minus signs mean significant exclusion of the topology, plus signs mean that the topology could not be excluded within the 95% confidence interval.

Table S14: **AU test results for the Platyhelminthes dataset with CAT-PMSF models.**

| Model | p-$AU_{T1}$ | | p-$AU_{T2}$ | |
|---|---|---|---|---|
| LG+CAT-PMSF chain 1 | 0.485 | + | $2.05 \cdot 10^{-6}$ | − |
| LG+CAT-PMSF chain 2 | 0.493 | + | $3.65 \cdot 10^{-5}$ | − |
| GTR+CAT-PMSF chain 1 | 0.497 | + | $7.44 \cdot 10^{-6}$ | − |
| GTR+CAT-PMSF chain 2 | 0.493 | + | $8.26 \cdot 10^{-3}$ | − |

T1 tree constraint is the same topology as the ML tree outputted by the CAT-PMSF pipeline, whereas T2 tree constraint is the ML tree under the specified model with a constraint producing the long branch attraction artifact. Minus signs mean significant exclusion of the topology, plus signs mean that the topology could not be excluded within the 95% confidence interval.

Table S15: **AU test results for the Simion et al. dataset with reduced and all the outgroups using CAT-PMSF models.**

| Dataset | Model | p-$AU_{T1}$ | | p-$AU_{T2}$ | |
|---|---|---|---|---|---|
| Simion reduced outgroup | LG+CAT-PMSF chain 1 | 0.606 | + | $7.73 \cdot 10^{-4}$ | − |
| Simion reduced outgroup | LG+CAT-PMSF chain 2 | 0.605 | + | $4.42 \cdot 10^{-4}$ | − |
| Simion reduced outgroup | GTR+CAT-PMSF chain 1 | 0.517 | + | $6.61 \cdot 10^{-4}$ | − |
| Simion reduced outgroup | GTR+CAT-PMSF chain 2 | 0.502 | + | $3.11 \cdot 10^{-4}$ | − |
| Simion all outgroups | LG+CAT-PMSF chain 1 | 0.604 | + | 0.396 | + |
| Simion all outgroups | LG+CAT-PMSF chain 2 | 0.587 | + | 0.413 | + |
| Simion all outgroups | GTR+CAT-PMSF chain 1 | 0.96 | + | $4.04 \cdot 10^{-2}$ | − |
| Simion all outgroups | GTR+CAT-PMSF chain 2 | 0.96 | + | $3.97 \cdot 10^{-2}$ | − |

T1 tree constraint is the same topology as the ML tree outputted by the CAT-PMSF pipeline, whereas T2 tree constraint is the ML tree under the specified model with a constraint producing the long branch attraction artifact. Minus signs mean significant exclusion of the topology, plus signs mean that the topology could not be excluded within the 95% confidence interval.

#### S6 Filtering out distant outgroups helps reduce long branch attraction

AU tests (Table S15) for the full Simion dataset (including 7 further outgroup species) could not reject the Ctenophora topology for LG+CAT-PMSF and barely rejected it for GTR+CAT-PMSF. This behavior is shown on the compositional constraint analysis (Fig. S16) where the CAT-PMSF results were not consistent, resembling the features of site-homogeneous and not complex enough site-heterogeneous models. Keeping only the closest outgroup, Choanoflagellatea (Fig. 4, S10 and S12), the AU test rejects the Ctenophora-sister topology (Table S15) and the CAT-PMSF model provides a weak, but consistent signal towards the Porifera-sister topology, which — based on the other cases presented in the main text — may imply that the Porifera-sister topology is the more probable animal root according to this dataset.

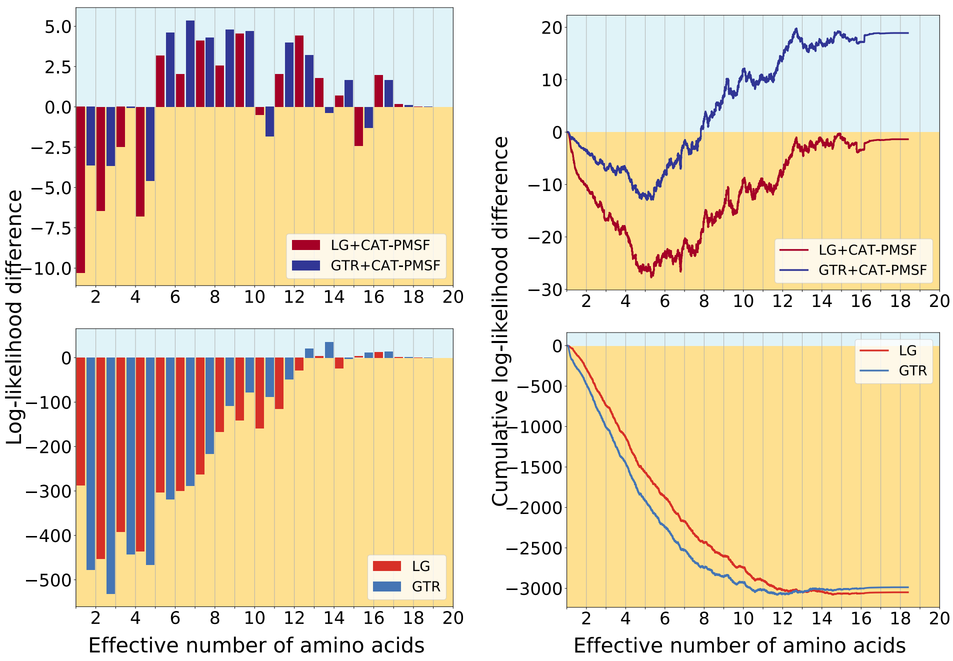

Figure S16: **Compositional constraint analysis of Simion et al.'s full outgroup dataset, binned and cumulative representation.** We performed analyses with the LG+CAT-PMSF, GTR+CAT-PMSF, LG (Le and Gascuel, 2008) and GTR (Tavaré, 1986) models constrained to either one of two competing topologies (the blue background supports the topology in Figure S5a, the yellowish background the topology in Figure S5b) with IQ-TREE 2 (Minh et al., 2020). On the left side the site-specific log-likelihood differences $\Delta\log L$ between the maximum likelihood trees of the two competing topologies binned according to their effective number of amino acids $K_{\text{eff}}$ estimated by PhyloBayes (Lartillot and Philippe, 2004) are shown. On the right side the cumulative site-specific log-likelihood differences $\Delta\log L$ between the maximum likelihood trees of the two competing topologies as a function of the sites with increasing effective number of amino acids $K_{\text{eff}}$ estimated by PhyloBayes (Lartillot and Philippe, 2004) are shown. Site-homogeneous models strongly favor Ctenophora at the animal root. LG+CAT-PMSF favors Ctenophora, while GTR+CAT-PMSF Porifera at the animal root, their signal is inconsistent.

#### S7 Results for another Metazoa dataset

We investigated a second dataset (Ryan et al., 2013) covering Metazoa. The results of the CAT-PMSF method suggest there is hardly any signal in the alignment discriminating between the two topologies (Fig. S17).

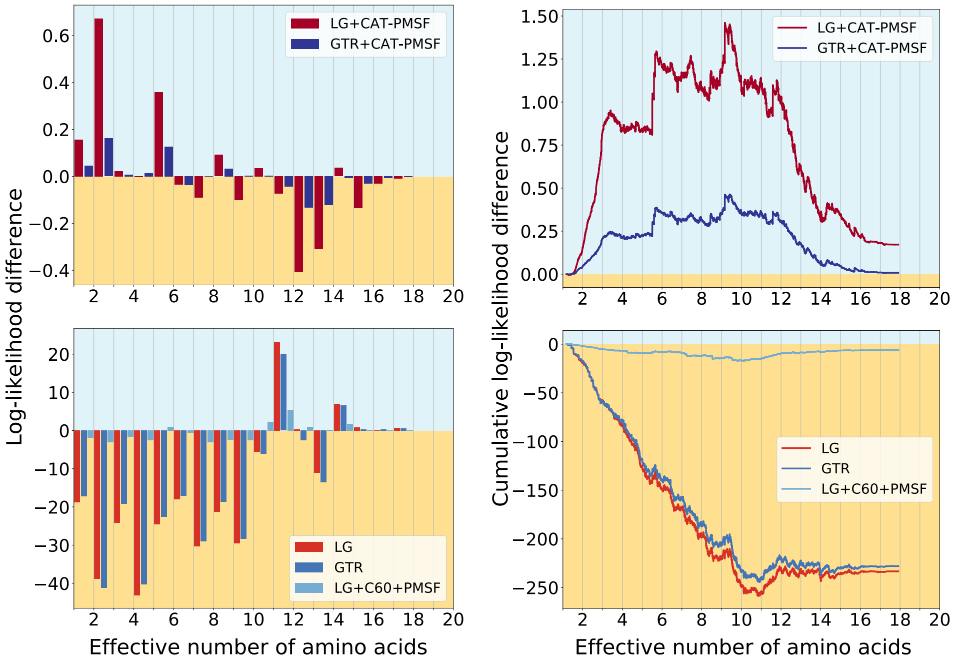

Figure S17: **Compositional constraint analysis of the Ryan et al. dataset, binned and cumulative representation.** We performed analyses with the LG+CAT-PMSF, GTR+CAT-PMSF, LG (Le and Gascuel, 2008), GTR (Tavaré, 1986) and LG+C60+PMSF (Quang et al., 2008; Wang et al., 2018) models constrained to either one of two competing topologies (the blue background supports the topology in Figure S6a, the yellowish background the topology in Figure S6b) with IQ-TREE 2 (Minh et al., 2020). On the left side the site-specific log-likelihood differences $\Delta \log L$ between the maximum likelihood trees of the two competing topologies binned according to their effective number of amino acids $K_{\mathrm{eff}}$ estimated by PhyloBayes (Lartillot and Philippe, 2004) are shown. On the right side the cumulative site-specific log-likelihood differences $\Delta \log L$ between the maximum likelihood trees of the two competing topologies as a function of the sites with increasing effective number of amino acids $K_{\mathrm{eff}}$ estimated by PhyloBayes (Lartillot and Philippe, 2004) are shown. Site-homogeneous models and the site-heterogeneous LG+C60+PMSF models show favor Ctenophora at the animal root. CAT-PMSF does not provide significant support to either of the topologies, there is not enough signal in the dataset.

#### S8 Compositional heterogeneity across sequences

CAT-PMSF assumes homogeneity of the evolutionary process across the branches of the tree. The software Homo v2.1 https://github.com/lsjermiin/Homo.v2.1 tests for compositional heterogeneity between sequences of an alignment. The simulated alignments do not show any signs of compositional heterogeneity between the sequences (for detailed results, see in the Dryad Digital Repository: https://doi.org/10.5061/dryad.g79cnp5rh and at https://github.com/drenal/cat-pmsf-paper). As expected, empirical alignments exhibit some level of compositional heterogeneity across branches. The following listings show the output of Homo v2.1 for the empirical datasets used in this study.

We observed that the significance of the test depends on the length of the sequences. In particular, we observed longer sequences reject compositional homogeneity with more significant p-values than shorter sequences (for detailed results, please see in the Dryad Digital Repository: https://doi.org/10.5061/dryad.g79cnp5rh and at https://github.com/drenal/cat-pmsf-paper). For example, the metazoan dataset of Ryan et al. (2013) has only 88k sites compared to approximately 400k sites of the metazoan dataset of Simion et al. (2017).

```
   Positions in alignment ....................... 24294
   Number of tests .............................. 780
   Smallest P value ............................. 2.400863e-256
   Level of significance (tau) .................. 0.050000
   Proportion of P values below tau ............. 0.778205
   Tests rejected (Bonferroni 1936) ............. 426
   Tests rejected (Holm 1979) ................... 434
   Tests rejected (Benjamini & Yekutieli 2001) .. 514
   Min(delta_Bowker) ............................ 0.006442
   Max(delta_Bowker) ............................ 0.021476
   Min(delta_EuclideanFS) ....................... 0.002527
   Max(delta_EuclideanFS) ....................... 0.023086
   Min(delta_EuclideanMS) ....................... 0.003250
   Max(delta_EuclideanMS) ....................... 0.069037
   Min(p distance) .............................. 0.094614
   Max(p distance) .............................. 0.603223
```

Listing S1: Results of Homo v2.1 for the dataset including Microsporidia.

```
Positions in alignment ...................... 35371
Number of tests ............................. 666
Smallest P value ............................ 1.002749e-37
Level of significance (tau) ................. 0.050000
Proportion of P values below tau ............ 0.681682
Tests rejected (Bonferroni 1936) ............ 300
Tests rejected (Holm 1979) .................. 309
Tests rejected (Benjamini & Yekutieli 2001) .. 370
Min(delta_Bowker) ........................... 0.005438
Max(delta_Bowker) ........................... 0.018051
Min(delta_EuclideanFS) ...................... 0.000995
Max(delta_EuclideanFS) ...................... 0.011907
Min(delta_EuclideanMS) ...................... 0.000941
Max(delta_EuclideanMS) ...................... 0.025573
Min(p distance) ............................. 0.029410
Max(p distance) ............................. 0.398590
```

Listing S2: Results of Homo v2.1 for the dataset including Nematoda.

```
Positions in alignment ...................... 35371
Number of tests ............................. 496
Smallest P value ............................ 1.139181e-38
Level of significance (tau) ................. 0.050000
Proportion of P values below tau ............ 0.675403
Tests rejected (Bonferroni 1936) ............ 232
Tests rejected (Holm 1979) .................. 239
Tests rejected (Benjamini & Yekutieli 2001) .. 283
Min(delta_Bowker) ........................... 0.005438
Max(delta_Bowker) ........................... 0.020186
Min(delta_EuclideanFS) ...................... 0.001709
Max(delta_EuclideanFS) ...................... 0.015175
Min(delta_EuclideanMS) ...................... 0.002571
Max(delta_EuclideanMS) ...................... 0.032438
Min(p distance) ............................. 0.047276
Max(p distance) ............................. 0.409459
```

Listing S3: Results of Homo v2.1 for the dataset including Platyhelminthes.

```
Positions in alignment ...................... 401632
Number of tests ............................. 4656
Smallest P value ............................ 0.000000e+00
Level of significance (tau) ................. 0.050000
Proportion of P values below tau ............ 0.987328
Tests rejected (Bonferroni 1936) ............ 4387
Tests rejected (Holm 1979) .................. 4470
Tests rejected (Benjamini & Yekutieli 2001) .. 4537
Min(delta_Bowker) ........................... 0.001975
Max(delta_Bowker) ........................... 0.012568
Min(delta_EuclideanFS) ...................... 0.000498
Max(delta_EuclideanFS) ...................... 0.015231
Min(delta_EuclideanMS) ...................... 0.000839
Max(delta_EuclideanMS) ...................... 0.042335
Min(p distance) ............................. 0.047812
Max(p distance) ............................. 0.441469
```

Listing S4: Results of Homo v2.1 for the metazoan dataset of Simion et al. (2017).

```
Positions in alignment ...................... 88384
Number of tests ............................. 1830
Smallest P value ............................ 6.090868e-255
Level of significance (tau) ................. 0.050000
Proportion of P values below tau ............ 0.362295
Tests rejected (Bonferroni 1936) ............ 289
Tests rejected (Holm 1979) .................. 289
Tests rejected (Benjamini & Yekutieli 2001) .. 392
Min(delta_Bowker) ........................... 0.000000
Max(delta_Bowker) ........................... 0.126419
Min(delta_EuclideanFS) ...................... 0.001647
Max(delta_EuclideanFS) ...................... 0.071180
Min(delta_EuclideanMS) ...................... 0.002744
Max(delta_EuclideanMS) ...................... 0.117840
Min(p distance) ............................. 0.059097
Max(p distance) ............................. 0.544691
```

Listing S5: Results of Homo v2.1 for the metazoan dataset of Ryan et al. (2013).

#### S9 Convergence measures of Phylobayes analyses

Here we show the output of Phylobayes' (Lartillot and Philippe, 2004) tool `tracecomp` to assess the convergence between the two Markov chains started for each alignment.

Table S16: **Convergence of Phylobayes LG+CAT+G4 analyses for the Microsporidia dataset**

| Parameter | Effective sample size | Relative difference |
|---|---|---|
| loglik | 594 | 0.0929612 |
| length | 596 | 0.0690003 |
| alpha | 231 | 0.129453 |
| Nmode | 623 | 0.0078885 |
| statent | 303 | 0.111053 |
| statalpha | 520 | 0.0183449 |

Two Markov chains were run, burnin for `tracecomp` was set to 4000.

Table S17: **Convergence of Phylobayes GTR+CAT+G4 analyses for the Microsporidia dataset**

| Parameter | Effective sample size | Relative difference |
|---|---|---|
| loglik | 766 | 0.0619622 |
| length | 180 | 0.171073 |
| alpha | 143 | 0.233501 |
| Nmode | 345 | 0.0253067 |
| statent | 242 | 0.0190515 |
| statalpha | 145 | 0.131871 |
| rrent | 282 | 0.0766182 |
| rrmean | 4773 | 0.00114388 |

Two Markov chains were run, burnin for `tracecomp` was set to 1000.

Table S18: **Convergence of Phylobayes LG+CAT+G4 analyses for the Nematoda dataset**

| Parameter | Effective sample size | Relative difference |
|---|---|---|
| loglik | 515 | 0.0539381 |
| length | 1296 | 0.0732629 |
| alpha | 798 | 0.0325119 |
| Nmode | 823 | 0.000358915 |
| statent | 719 | 0.0628802 |
| statalpha | 316 | 0.0598403 |

Two Markov chains were run, burnin for `tracecomp` was set to 1000.

Table S19: **Convergence of Phylobayes GTR+CAT+G4 analyses for the Nematoda dataset**

| Parameter | Effective sample size | Relative difference |
|---|---|---|
| loglik | 540 | 0.15339 |
| length | 5339 | 0.250924 |
| alpha | 4724 | 0.182099 |
| Nmode | 1292 | 0.0321017 |
| statent | 301 | 0.00237055 |
| statalpha | 419 | 0.180082 |
| rrent | 342 | 0.130712 |
| rrmean | 6179 | 0.00141907 |

Two Markov chains were run, burnin for `tracecomp` was set to 1000.

Table S20: **Convergence of Phylobayes LG+CAT+G4 analyses for the Platyhelminthes dataset**

| Parameter | Effective sample size | Relative difference |
|---|---|---|
| loglik | 110 | 0.0514937 |
| length | 634 | 0.04558 |
| alpha | 568 | 0.0253402 |
| Nmode | 3691 | 0.0366779 |
| statent | 137 | 0.0540562 |
| statalpha | 166 | 0.0345426 |

Two Markov chains were run, burnin for `tracecomp` was set to 1000.

Table S21: **Convergence of Phylobayes GTR+CAT+G4 analyses for the Platyhelminthes dataset**

| Parameter | Effective sample size | Relative difference |
|---|---|---|
| loglik | 359 | 0.351519 |
| length | 418 | 0.0778552 |
| alpha | 401 | 0.43393 |
| Nmode | 347 | 0.0557145 |
| statent | 381 | 0.262808 |
| statalpha | 430 | 0.102403 |
| rrent | 2906 | 0.117868 |
| rrmean | 7249 | 0.0108546 |

Two Markov chains were run, burnin for `tracecomp` was set to 1000.

Table S22: **Convergence of Phylobayes LG+CAT+G4 analyses for the Simion et al. dataset with Choanoflagellatea as outgroup**

| Parameter | Effective sample size | Relative difference |
|---|---|---|
| loglik | 147 | 0.381065 |
| length | 41 | 0.129443 |
| alpha | 151 | 0.323356 |
| Nmode | 218 | 0.107418 |
| statent | 139 | 0.180409 |
| statalpha | 72 | 0.0778117 |

Two Markov chains were run, burnin for `tracecomp` was set to 1000.

Table S23: **Convergence of Phylobayes GTR+CAT+G4 analyses for the Simion et al. dataset with Choanoflagellatea as outgroup**

| Parameter | Effective sample size | Relative difference |
|---|---|---|
| loglik | 159 | 0.480894 |
| length | 107 | 1.19047 |
| alpha | 87 | 1.94857 |
| Nmode | 115 | 0.168816 |
| statent | 111 | 3.54829 |
| statalpha | 48 | 0.370279 |
| rrent | 136 | 0.734394 |
| rrmean | 2461 | 0.0347738 |

Two Markov chains were run, burnin for `tracecomp` was set to 1000.

Table S24: **Convergence of Phylobayes Poisson+CAT+G4 analyses for the Simion et al. dataset with Choanoflagellatea as outgroup**

| Parameter | Effective sample size | Relative difference |
|---|---|---|
| loglik | 142 | 0.0572755 |
| length | 1009 | 0.0122319 |
| alpha | 661 | 0.0971534 |
| Nmode | 118 | 0.106781 |
| statent | 282 | 0.0674665 |
| statalpha | 172 | 0.0401437 |

Two Markov chains were run, burnin for `tracecomp` was set to 3000.

Table S25: **Convergence of Phylobayes LG+CAT+G4 analyses for the Simion et al. dataset including distant outgroups**

| Parameter | Effective sample size | Relative difference |
|---|---|---|
| loglik | 199 | 0.0207354 |
| length | 478 | 0.000169223 |
| alpha | 559 | 0.0570139 |
| Nmode | 333 | 0.0671745 |
| statent | 591 | 0.0899417 |
| statalpha | 117 | 0.151863 |

Two Markov chains were run, burnin for `tracecomp` was set to 1000.

Table S26: **Convergence of Phylobayes GTR+CAT+G4 analyses for the Simion et al. dataset including distant outgroups**

| Parameter | Effective sample size | Relative difference |
|---|---|---|
| loglik | 33 | 1.92869 |
| length | 261 | 0.124522 |
| alpha | 94 | 2.27226 |
| Nmode | 179 | 0.22674 |
| statent | 74 | 2.6127 |
| statalpha | 204 | 0.0657509 |
| rrent | 205 | 0.339186 |
| rrmean | 4999 | 0.003295 |

Two Markov chains were run, burnin for `tracecomp` was set to 1000.

Table S27: **Convergence of Phylobayes LG+CAT+G4 analyses for the Ryan et al. dataset**

| Parameter | Effective sample size | Relative difference |
|---|---|---|
| loglik | 313 | 0.0435181 |
| length | 227 | 0.0927871 |
| alpha | 167 | 0.052443 |
| Nmode | 327 | 0.00677682 |
| statent | 297 | 0.0662057 |
| statalpha | 274 | 0.0772407 |

Two Markov chains were run, burnin for `tracecomp` was set to 3000.

Table S28: **Convergence of Phylobayes GTR+CAT+G4 analyses for the Ryan et al. dataset**

| Parameter | Effective sample size | Relative difference |
|---|---|---|
| loglik | 294 | 0.176161 |
| length | 626 | 0.0480805 |
| alpha | 292 | 0.0418603 |
| Nmode | 681 | 0.225009 |
| statent | 136 | 0.176489 |
| statalpha | 64 | 0.0754489 |
| rrent | 265 | 0.025304 |
| rrmean | 2861 | 0.0100864 |

Two Markov chains were run, burnin for `tracecomp` was set to 2700.
